# Amino acid restriction sensitizes lung cancer cells to ferroptosis via GCN2-dependent activation of the integrated stress response

**DOI:** 10.1101/2025.04.10.648120

**Authors:** Viktor J. Garellick, Nadia Gul, Parvin Horrieh, Dyar Mustafa, Angana A.H. Patel, Martin Dankis, Samantha Alvarez, Johanna Berndtson, Maria Schwartz, Andreas Persson, Fikret Zahirovic, Clotilde Wiel, Volkan I. Sayin, Per Lindahl

**Affiliations:** Sahlgrenska Center for Cancer Research, University of Gothenburg, 405 30 Gothenburg, Sweden; Department of Molecular and Clinical Medicine, Institute of medicine, University of Gothenburg, 405 30 Gothenburg, Sweden; Department of surgery, Institute of clinical medicine, University of Gothenburg, 405 30 Gothenburg, Sweden; The Wallenberg Centre for Molecular and Translational Medicine, University of Gothenburg, 405 30 Gothenburg, Sweden; Department of medical biochemistry and cell biology, Institute of Biomedicine, at Sahlgrenska Academy, University of Gothenburg, 405 30 Gothenburg, Sweden; Department of Nutritional Physiology, Institute of Nutritional Sciences, Friedrich Schiller University Jena, Dornburger Str. 24, 07743, Jena, Germany

**Keywords:** lung cancer, amino acids, glutathione, ferroptosis, integrated stress response, mitochondrial respiration

## Abstract

Lung cancer cells are vulnerable to iron-dependent oxidation of phospholipids leading to ferroptosis, a process countered by glutathione peroxidase-4 that converts lipid hydroperoxides to lipid alcohols using glutathione as reducing agent. Since ferroptosis-inducing agents are in clinical development, identifying modifiers of ferroptosis susceptibility is warranted. Here, we investigate the impact of amino acids on susceptibility to buthionine sulfoximine (BSO), a glutamate-cysteine ligase inhibitor that blocks biosynthesis of glutathione. We found that reduced amounts of amino acids other than cysteine increased the sensitivity to BSO and other ferroptosis-inducing agents, in a panel of mouse and human lung cancer cells, without affecting glutathione production. Activation of the amino acid sensor protein GCN2 and the integrated stress response lowered the threshold for lipid peroxidation by stimulating ATF4-dependent mitochondrial respiration. The finding has implications for lung cancer metabolism and raises the possibility of using protein restricted diets in combination with ferroptosis-inducing agents as cancer therapies.

## 1. Introduction

The cellular redox environment is firmly controlled and deviations from homeostasis may cause oxidative stress, a state where the amount of reactive oxygen species (ROS) exceeds the antioxidant capacity. Oxidative stress has emerged as a limiting factor for cancer progression and metastasis. Lung cancer cells are particularly sensitive to oxidative stress, as evidenced by increased lung cancer risk in clinical trials of dietary antioxidants [1, 2], the accumulation of somatic mutations in nuclear factor erythroid 2-related factor 2 (NFE2L2, NRF2) or kelch-like ECH-associated protein 1 (KEAP1) in lung cancer cells [3–5], and mechanistic studies in mice [6, 7]. Prooxidant cancer therapies that target endogenous antioxidants may therefore be effective against lung cancer.

Glutathione is the most abundant intracellular antioxidant and a major determinant of the cellular redox environment [8]. One important role of glutathione is to maintain the integrity of phospholipid membranes. Glutathione peroxidase-4 (GPX4) plays a key role in this process by converting harmful lipid hydroperoxides to benign lipid alcohols, using glutathione as reducing agent [9]. Depletion of glutathione leads to ferroptosis, a regulated form of cell death that is triggered by iron-dependent lipid peroxidation [10]. Ferroptosis has emerged as a central non-apoptotic cell death mechanism in cancer, and ferroptosis-inducing agents are considered for clinical development [11]. Identifying factors that govern susceptibility to ferroptosis-inducing agents could lead to improved therapeutic options.

Cultured human cancer cells are surprisingly resilient to glutathione depletion [12–14]. This can be explained by compensatory upregulation of the thioredoxin system [15–18], but other factors may contribute. Tumor angiogenesis produces abnormal blood vessels leading to hypoxia and nutrient stress in tumors. Results obtained with culture media designed to optimize cell growth, such as the aforementioned studies, may underestimate the impact of nutrient stress on ferroptosis.

Ferroptosis susceptibility is modified by lipid metabolism, iron homeostasis, and amino acid metabolism, the latter mainly by influencing levels of glutathione [11]. The tripeptide glutathione is synthesized from glutamate, cysteine, and glycine in reactions that are limited by cysteine availability, and is hence vulnerable to perturbations affecting cysteine metabolism [19, 20]. However, amino acids are major modifiers of drug resistance that intersect with drug mechanisms in abundant and variable fashions [21, 22]. Whether amino acids modify ferroptosis susceptibility in additional ways beyond glutathione synthesis warrants further investigation.

Here, we investigate the impact of levels of amino acids on sensitivity to buthionine sulfoximine (BSO), a glutamate-cysteine ligase catalytic subunit (GCLC) inhibitor that irreversibly blocks *de novo* biosynthesis of glutathione [23]. GCLC catalyzes the condensation of cysteine and glutamate and is the rate limiting enzyme in glutathione biosynthesis [24].

## 2. Materials and methods

### 2.1 Cell culture

The cell lines A549, NCIH838 (H838), NCIH1299 (H1299), NCIH23 (H23), NCIH460 (H460), SKNEB2, and SHSY5Y were obtained from ATCC. Cas9-expressing A549 cells were obtained from GeneCopoeia (SL-504; GeneCopoeia, Inc., Rockville, MD) and the mouse KP cell line was obtained from V. Sayin and is described [25].

A549, NCIH838, NCIH1299, NCIH23, and NCIH460 cells were maintained in Ham’s F12 mixture (SH30026.01; Hyclone) or RPMI1640 (lot nr RB35958, cat nr SH30027.01; Hyclone), supplemented with 1% penicillin/streptomycin (SV30010; Hyclone) and 10% FBS (SV30160.03; Hyclone). SKNEB2 and SHSY5Y were maintained in F12:MEM (1:1) supplemented with 10% FBS and 1% penicillin/streptomycin. KP cells were maintained in DMEM with 10% FBS and 0.1% gentamicin (15710049; Gibco). Experiments on KP cells were carried out in F12 or RPMI supplemented with 10% FBS and 0,1% gentamicin.

Custom-made F12AA and RPMIAA (Genaxxon bioscience, Germany; see Table 1 for medium formulation) were supplemented with 35.15 mg/L final concentration of L-cysteine-hydrochloride-monohydrate (C6852; Sigma Aldrich), 146 mg/L final concentration of L-glutamine (25030-081; Gibco), 1% penicillin/streptomycin, and 10% FBS. Cells were switched to experimental conditions one passage prior to the experiment unless otherwise specified in the text. Cells were kept at 37°C and 5% CO2.

**Table 1.**
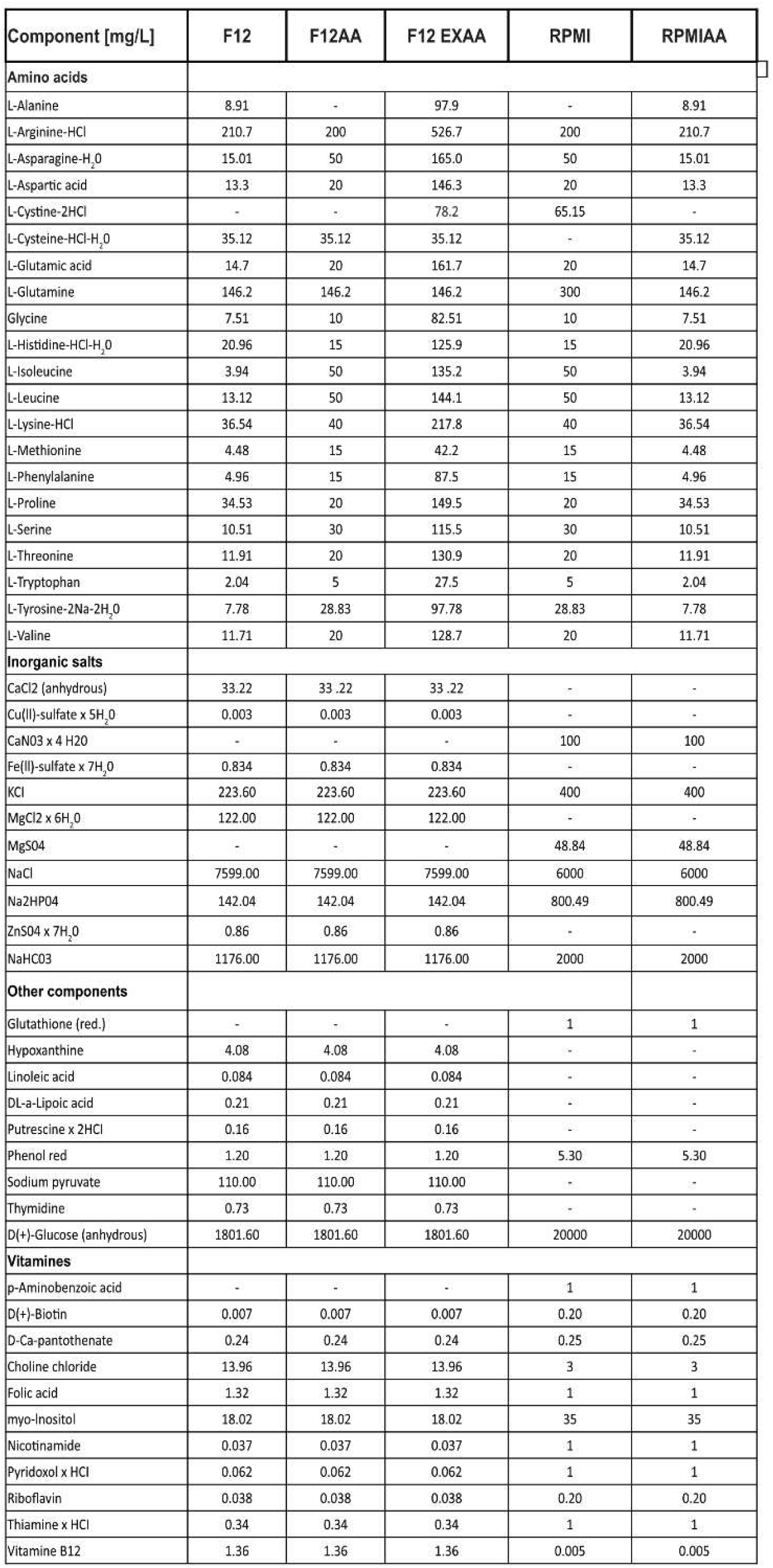

### 2.2 Cell growth

1.5x10^5^ cells/well were seeded in a 6-well plate in F12 or RPMI. On the first passage, after three days, the cells were counted using a trypan blue exclusion assay. 1.5x10^5^ cells/well were then re-seeded and counted again for two more passages.

### 2.3 7-AAD staining

1.5x10^6^ cells/well were seeded overnight in a 6-well plate. The next day, cells were incubated in 5μM etoposide overnight to induce apoptosis, after which the cells were washed in cold PBS and incubated with 1ng/µl 7-AAD (A1310; Thermo Fisher) for 30 minutes on ice. The 7AAD signal was quantified with a flow cytometer (BD Accuri C6) at (λex488nm) using the FL3 detector. The cut-off value for 7-AAD positive cells were based on the signal distribution in etoposide-treated cells versus untreated negative controls.

### 2.4 Cell viability

For crystal violet viability assays, 2x10^4^ cells/well were seeded overnight in 12-well plates, incubated with drugs dissolved in fresh media for 72 hours, and stained with 0,025% (v/v) of crystal violet staining solution (V5265; Sigma Aldrich) supplemented with 1% (v/v) formaldehyde, 1% (v/v) methanol and diluted in PBS, for 20 minutes. After staining, the plates were washed with tap water and air dried for a minimum of 24 hours. Once dried, the staining solution was re-dissolved with 10% acetic acid and light absorbance were measured at 595 nm using a SpectraMax MiniMax spectrophotometer. Absorbance was subtracted for background and then normalized to untreated control.

CellTiter-Glo® (G7570; Promega) assays were performed according to the manufacturer’s instruction. Briefly, 2x10^3^ cells/well were seeded overnight in white opaque-bottom 96-well plates, incubated with drugs in fresh medium for 72 hours, and stained with CellTiter-Glo® reagent for 10 minutes. Luminescence was measured with a BioTek Synergy HTX plate reader. Luminescence was normalized to untreated control.

### 2.4 Drug treatments

Drugs used were auranofin (A6733; Sigma-Aldrich), α-tocopherol (T3251; Sigma-Aldrich), bafilomycin A1 (SML1661; Sigma-Aldrich), certolizumab pegol (HYP9953; MedChemExpress), chloroquine (C6628; Sigma-Aldrich), CU-CPT4A (HY-108473; MedChemExpress), deferoxamine (D9533; Sigma-Aldrich), erastin (e7781;Sigma Aldrich), FCCP (Carbonyl cyanide 4-(trifluoromethoxy)phenylhydrazone) (HY-100410; MedChemExpress), ferrostatin-1 (SML0583; Sigma-Aldrich), L-buthionine-sulfoximine (b2515; Sigma-Aldrich), necrostatin-1 (N9037; Sigma-Aldrich), liproxstatin (SML1414; Sigma-Aldrich), oligomycin (HY-N6782; MedChemExpress), puromycin dihydrochloride (A1113803;Gibco), rotenone (HY-B1756; MedChemExpress), RSL3 (SML2234; Sigma-Aldrich), resatorvid (HY-11109; MedChemExpress), and Z-VAD-FMK (V116; Sigma-Aldrich).

### 2.6 Western blotting

Semi-confluent cells were lysed with 8M Urea buffer or RIPA buffer supplemented with Pierce protease and phosphatase inhibitor cocktail (A32959; Thermo Fisher Scientific), centrifuged at 20000 rcf for 10 minutes at 4°C, mixed with LDS loading buffer supplemented with 10mM DTT or 2.5% β-mercaptoethanol, and incubated at 70°C for 10 minutes. Proteins were separated on SDS-PAGE gels at 200V and transferred onto nitrocellulose membranes using semi-dry transfer. Membranes were blocked with EveryBlot blocking buffer (12010020; Bio-Rad) for 5 minutes, or with 5% milk in TBS-T buffer for 60 minutes. Primary antibody incubation was done overnight in blocking buffer at 4°C and secondary incubation at room temperature for 1 hour. Membranes were incubated with Clarity™ Western ECL substrate (1705060; Bio-Rad) for 5 minutes. Chemiluminescent imaging was done using a GE Amersham Imaging device. Band intensities were quantified with the ImageJ software. Primary antibodies used were 4E-BP1 (9452; Cell signaling technology (CST)), ATF4 (11815; CST), α-Tubulin (T6199; Sigma-Aldrich), CHAC1 (15207-1-AP;Proteintech), CHOP (15204-1; Proteintech), eIF2α (5324; CST), Ferritin (ab75973-1001; Abcam), GADD34 (10449-1-AP;Proteintech), GAPDH (5174; CST), GCN2 (3302; CST), GCLC (ab41463; Abcam), LC3B (NB100-2220SS; Novusbio), p-4E-BP1 (2855; CST), p-eIF2α Ser 51 (3398; CST), p-GCN2 (ab75836; Abcam), p-S6 (2211; CST), S6 (2217; CST), TfrC (13-6800;Invitrogen), p-PKR (ab32036; Abcam), p-PERK (ab192591; Abcam), PERK (3192; CST), PKR (sc-6282; SCBT), and Hsp90 (4877; CST). Secondary antibody used was peroxidase-conjugated AffiniPure Goat Anti-Rabbit IgG (H+L) (111-035-003; Jackson Immunoresearch) and peroxidase-conjugated AffiniPure Goat Anti-Mouse IgG (H+L) (115-035-003; Jackson Immunoresearch).

### 2.7 qPCR

Total RNA was isolated from A549 cells transfected with siRNA 24-hours post-transfection, using RNeasy mini kit (Qiagen) according to the manufacturer’s protocol. The purity and concentration of the extracted RNA were assessed using a NanoDrop spectrophotometer. cDNA was synthesized from 300 ng of total RNA using the iScript™ Reverse Transcription Supermix for RT-qPCR (Biorad) according to the manufacturer’s protocol. The reverse transcription reaction was carried out in a thermal cycler at 25°C for 5 minutes, followed by 46°C for 20 minutes, and then 95°C for 1 minutes to inactivate the enzyme. qPCR reactions were performed in a 20 μl volume containing 2 μl of cDNA, 10 μl of PowerUp™ SYBR™ Green Master Mix (Thermo Fisher), 300nM of each primer, and nuclease-free water to a final volume of 20 μl. Primer pairs against ASNS, SLC7A11, CHAC1 and DDIT3/CHOP were purchased from Sigma-Aldrich (KiCqStart™ Primers ID ASNS ID H_ASNS_1, SLC7A11 ID H_SLC7A11_1, DDIT3/CHOP H_DDIT3_1, CHAC1 H_CHAC1_1). Primers pairs against GAPDH was purchased from IDT technologies (Forward primer 5’-CGC TCT CTG CTC CTC CTG TT-3’ and Reverse primer 5’-CCA TGG TGT CTG AGC GAT GT-3’). The qPCR was carried out in an Applied Biosystems 7900 Real-Time PCR System with cycling conditions as per manufacturer’s recommendations. Gene expression levels were normalized to GAPDH and analyzed using the 2^-ΔΔCt method. No-template controls were included in each run, and melting curve analysis was performed to ensure specificity of the PCR products.

### 2.8 Generation of GCLC knockout cells

GCLC knockout cells were generated with CRISPR-Cas9, as in [26]. Briefly, guide-RNA (gRNA) spacer sequences targeting constituently expressed exons in GCLC were designed using the Benchling tool (https://www.benchling.com). Double-stranded DNA oligonucleotides (20nt) were cloned into the BamH1 site of the Lentiguide puro vector, and co-transfected with pCMV-R8.2 and pCMV-VSV-G into HEK293T cells to produce gRNA coding lentivirus. Cas9-expressing A549 cells were infected with virus and cultured in DMEM supplemented with puromycin for 7 days to produce knockout batch clones (verified with western blotting at day 7; **Fig. S1F**) that were used in experiments. The cells were switched to F12 after day 7 and evaluated for viability at day 13. Lentiguide puro was a kind gift from Feng Zhang (Addgene plasmid #52963; http://n2t.net/addgene: 52963; RRID:Addgene_52963) [27]. Oligonucleotide sequences used for the creation of gRNA constructs were: GCLC gRNA#1: forward oligo CACCGCATACTCACCTGAAGCGA, reverse oligo AAACTCGCTTCAGGTGAGTATGC; GCLC gRNA#2: forward oligo CACCGTAGATGTGCAGGAACTGG, reverse oligo AAACCCAGTTCCTGCACATCTAC; GCLC gRNA#3: forward oligo CACCGAAATATCCGACATAGGAG, reverse oligo AAACCTCCTATGTCGGATATTTC; control gRNA: forward oligo CACCGGAGGCTAAGCGTCGCAA, reverse oligo AAACTTGCGACGCTTAGCCTCC.

### 2.8 CM-H2DCFDA ROS measurements

For estimation of general ROS levels, A549 cells were seeded at a density of either 100,000 cells per well in a 6-well plate containing RPMI or F12 media and allowed to attach overnight. The cells were then treated with or without 0.1 mM BSO for 24 h. As positive controls, cells were treated for 10 min with 25 µM Menadione. Following treatments, the cells were harvested using trypsin and subsequently incubated for 30 min at 37°C with 5 µM CM-H2DCFDA solution (C6827; Thermo Fisher Scientific). The cells were then washed three times with PBS, resuspended in 200 µL of PBS, and flow cytometry data was collected and analyzed using a BD Accuri™ C6 Plus Flow Cytometer.

### 2.9 Lipid peroxidation

Lipid peroxidation was estimated using the lipid soluble redox-sensitive BODPIPY-C11™ sensor (D3861, Thermo Fisher) as in [28]. A549 or H838 cells were plated in 6-well plates at 1x10^5^ and 2x10^5^ cells/well, respectively, in the presence of 100µM BSO or vehicle. 24-hours post plating, BODIPY-C11 was added to the culture medium to a final concentration of 1.5μM, after which the cells were stained for 20 minutes at 37°C. The amount of oxidized BODIPY-C11 was quantified with a flow cytometer (BD Accuri C6) at (λex488nm) using the FL1 detector. To define BODIFY-C11 positive cells, a cut-off value was calculated based on the signal distribution in positive (cells treated with 5μM erastin for 6 hours) and negative (untreated cells) controls.

### 2.10 Glutathione determination

Glutathione (reduced) concentrations were determined using the GSH-Glo™ Glutathione assay (V6911; Promega corporation) according to the manufacturer’s instruction. 2000 cells/well were seeded overnight in white opaque-bottomed 96-well plates. The next day, cells were incubated with BSO (the concentrations are indicated in the graphs) in fresh medium for 24 hours, after which GSH-Glo and luciferin detection reagents were added. Luminescence was measured using a BioTek Synergy HTX plate reader.

### 2.11 Gas chromatography-mass spectrometry (GC/MS) analysis of amino acids

For the amino acid uptake assay, 1 x 10⁵ cells were seeded in 2 mL of their respective cell culture medium with 10% FBS in 6 well plates for two days before harvest. For relative determination of intracellular amino acid levels, 1x 10⁵ cells were seeded in 2 mL of RPMI and left to attach overnight. The next day, spent cell medium was replaced with fresh F12, F12AA and RPMI. Lysates were collected at 1, 6, 24 and 48 hours post medium renewal by scraping the cells in 200 µl of 80% (v/v) ice cold methanol containing 1 µg/mL norvaline (Sigma-Aldrich). All Samples were then vortexed for 10 min at 4°C and then centrifuged at 20 000 rcf for 10 min. The supernatant was transferred to fresh tubes and solvents were evaporated in a speed vac for 2 h. Dried metabolite extracts were subsequently derivatized with 20 µL O-methoxyamine-hydrochloride (MOX) reagent (Sigma-Aldrich) in pyridine (Sigma-Aldrich) at a concentration of 20 mg/mL for 60 min at 37°C followed by 20 µL of N-tert-butyldimethylsilyl-N-methyltrifluoracetamide with 1% tert-butyldimethylchlorosilane (TBDMS, Sigma-Aldrich) for 60 min at 37°C. After derivatization, samples were analyzed by GC-MS using an DB-35ms column (Agilent Technologies) in an Agilent Intuvo gas chromatograph coupled to an Agilent 5997B mass spectrometer. Helium was used as the carrier gas at a flow rate of 1.2 mL/min. 1 µl of sample was injected in split mode (split 1:1) at 270°C. After injection the GC oven was held at 100°C for one min and then increased to 300°C at 3.5°C/min. The oven was then ramped to 320°C at 20°C/min and held for 5 min at 320°C. The MS system operated under electron impact ionization at 70 eV and the MS source and quadrupole were held at 230°C and 150°C respectively. The detector was used in scanning mode, and the scanned ion range was 10-650 m/z. Mass isotopomer distributions were determined by integrating the appropriate ion fragments for each metabolite [29] using MATLAB (Mathworks) and an algorithm adapted from Fernandez and colleagues [30] that corrects for natural abundance. For the uptake assay, total metabolite pool sizes were normalized to cell counts for each condition separately. For intracellular amino acid determination, total metabolite pool sizes were normalized to total protein content determined by Pierce BCA assay for each condition separately.

### 2.12 Overexpression of CHOP and ATF4

To achieve overexpression of CHOP and ATF4, the plasmids TFORF0623 and TFORF3036 were used respectively. Both plasmids were obtained from Addgene (plasmid #142666 and plasmid #144512, respectively) as gifts from Feng Zhang. Lentiviral particles were produced (as described above) from these plasmids and used to transduce A549 cells. Following transduction, the culture medium was replaced after 24 hours and cells were selected in puromycin for 3 days to isolate successfully transduced populations. The selected cells were subsequently used in downstream experiments to assess the effects of CHOP and ATF4 overexpression.

### 2.12 siRNA transfection

2x10^5^ or 2x10^3^ cells/well were seeded overnight in 6- or 96-well plates, respectively, and then transfected with 25 or 1 pmol siRNA duplexes (or a universal negative control) using Lipofectamine ™ RNAiMax transfection reagent (13778100; Thermo Fisher Scientific), according to the manufacturer’s instruction. 24-48 hours after transfection, the cells were used in experiments or harvested for downstream analyses. siRNAs used were SASI_HS01_00097889, SASI_HS01_00097888, SASI_HS01_00097890, SASI_HS02_00332313, SASI_HS02_00332314, SASI_HS01_00175197, SASI_HS01_00153013, SASI_HS02_00336880 and SASI_HS01_00153015 (Sigma-Aldrich).

### 2.13 SeaHorse analysis of mitochondrial respiration

Analyses of mitochondrial respiration were performed using the Seahorse XFe96/XFPro Analyzer (Agilent), which measures the extracellular acidification rate (ECAR) and oxygen consumption rate (OCR) of live cells. Mitochondrial stress was measured using the following compounds: 0.5μM oligomycin, 1μM FCCP, and 0.5μM rotenone. Cells were seeded in Seahorse XFe96/XF Pro Cell Culture Microplates (10,000 cells/well) (103792-100, Agilent) and cultured overnight in F12 or F12AA medium at 37°C in a CO2 incubator. 45 minutes before the assay, the medium was replaced with freshly prepared phenol red–free RPMI medium (Agilent 103576-100) supplemented with 1 mM glucose, and 2 mM glutamine. Prior to the assays, the Seahorse Analyzer was pre-warmed to 37°C. ATP production rates were calculated using the Macro Excel file provided by the manufacturer.

### 2.14 Ratiometric confocal microscopy

Images were acquired from live cells on an LSM 980 (Zeiss) using a 63x/1.40 oil immersion plan-apochromatic objective lens kept at 37°C. A sequential acquisition was setup with the first track excitation at 561nm and detection from 552 to 640nm, imaging the oxidized BODIPY-C11, and the second track excitation at 488 and 639nm and detection from 493 to 517nm and 653 to 668nm, imaging the reduced BODIPY-C11 and the Mitotracker DeepRed (Invitrogen) or LipidSpot610 (Biotium), respectively. 7–16 random fields per condition were imaged using the ZEN software Sample Navigator for unbiased sampling.

### 2.15 Statistics

Statistical analyses were done with GraphPAD Prism 9 (GraphPad Software, San Diego, CA, USA). Difference between two groups was tested with unpaired student’s t-test, difference between multiple groups and treatments was tested with multiple unpaired t-test, one-way ANOVA or two-way ANOVA using Dunnet’s or Tukey’s post-hoc test (main row effect), and difference in LC3-II levels and cell growth was tested with multiple unpaired t-tests. Linear regression for autophagy flux was calculated with GraphPad Prism 9. IC50 values were calculated using non-linear regression analysis of inhibitor versus response: variable slope (four parameters). *P*-values <0.05 was considered significant. Data are presented as mean ± SEM.

## 3. Results

### 3.1 Nutritional factors influence the susceptibility to ferroptosis-inducing agents in cancer cells

To investigate the impact of nutrient factors on susceptibility to ferroptosis-inducing agents, we established cultures of the human lung cancer cell line A549 in Ham’s F12 nutrient mixture (F12) or RPMI, two culture media used in cancer research that differ in amino acid content. Culture in F12 retarded cell growth slightly compared to culture in RPMI (**Fig. S1A**) but had no effect on cell death or cell morphology (**Fig. S1B, C**). To assess the susceptibility to ferroptosis-inducing agents, we established dose response curves for BSO. While RPMI-cultured cells were strikingly resistant to BSO, F12-cultured cells died at low micromolar concentrations (**Fig. 1A, B**). Similar results were obtained for H838, H1299, and H23 lung cancer cells (**Fig. 1B**) indicating that nutrient factors have significant impact on BSO sensitivity.

**Fig. 1.**
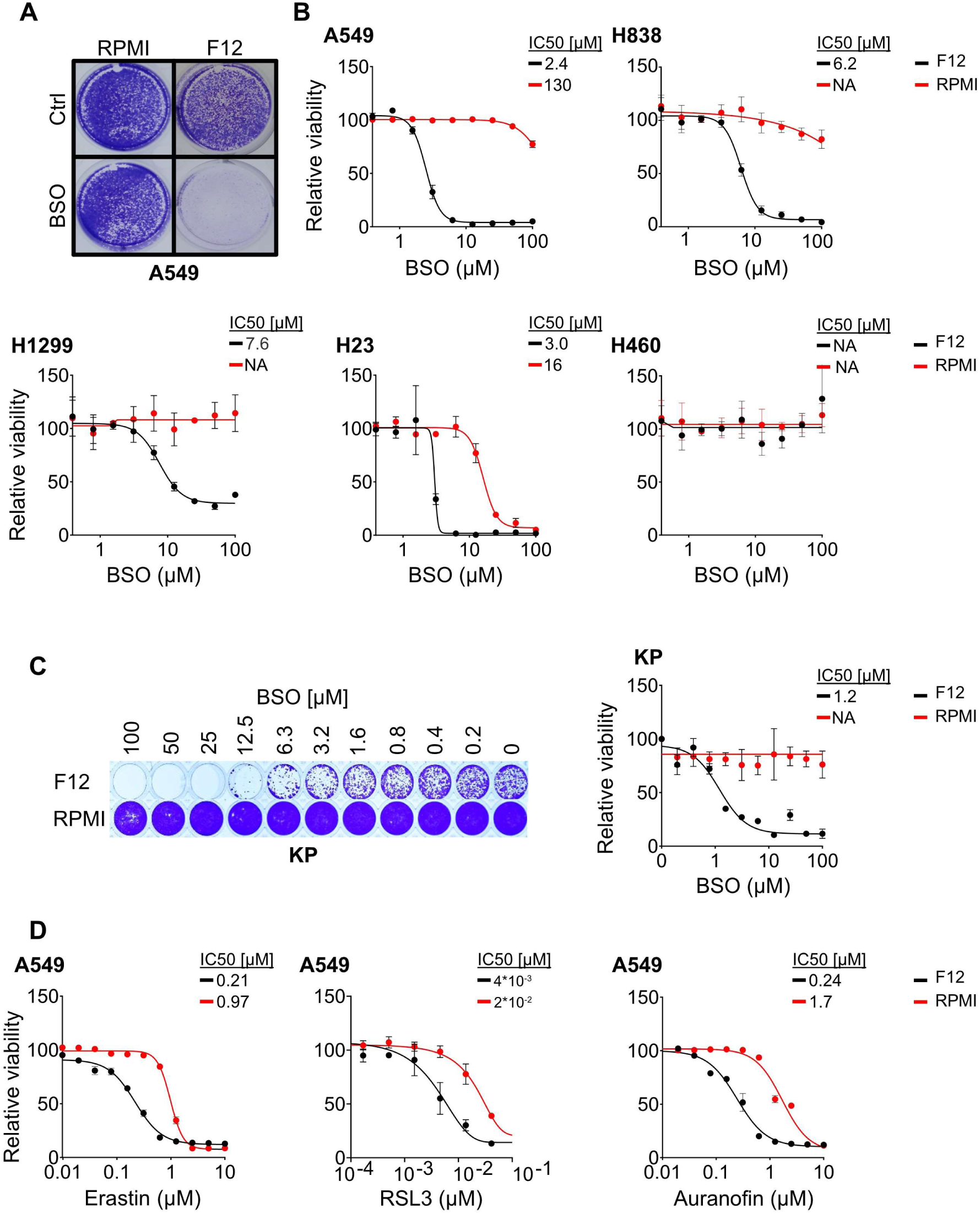
Culture in F12 medium sensitizes lung cancer cells to BSO. (**A**) Crystal violet staining of A549 cells that were cultured in RPMI or F12, after treatment with 100μM BSO or vehicle (Ctrl) for 72 hours. (**B**) BSO dose response curves for A549, H838, H1299, H23, and H460 cells cultured in RPMI or F12 for 72 hours. The data were normalized against the mean of the untreated samples for each condition. (**C**) Crystal violet staining and quantification of mouse KP cells that were cultured in RPMI or F12, in the presence of BSO at the indicated concentrations for 72 hours. (**D**) Dose response curves for A549 cells cultured in RPMI or F12 and treated with Erastin, RSL3, or Auranofin for 72 hours. The data were normalized against the mean of the untreated samples for each condition. n=3 replicates for all datapoints, error bars show SEM.

Culture in F12 increased BSO sensitivity in mouse KP lung cancer cells (**Fig. 1C**) and human SKNBE-2 neuroblastoma cells compared to RPMI (**Fig. S1D**), showing that the mechanism is evolutionarily conserved and not restricted to lung cancer cells. The culture media had no impact on BSO sensitivity in human H460 lung cancer cells or SH-SY5Y neuroblastoma cells (**Fig. 1B** and **Fig. S1D**), indicating that intrinsic factors such as genetic makeup, epigenetic history, or cell-of-origin play a role. However, a comparison of COSMIC Hot Spot mutations between BSO sensitive and insensitive cell lines showed no obvious relationship between drug responsiveness and genotype (**Fig. S1E**).

To verify that BSO-lethality is caused by GCLC-inhibition, A549 cells were infected with lentivirus expressing sgRNAs targeting GCLC, or a non-targeting control. Knockout of GCLC was lethal in F12-cultured cells, consistent with the pharmacological data (**Fig. S1F, G**).

To test whether F12 increases the sensitivity to other ferroptosis-inducing agents, we established dose response curves for erastin, RSL3 and auranofin in F12 or RPMI-cultured A549 cells. The IC50 values of the drugs were consistently lower in cells that were cultured in F12 compared to RPMI (**Fig. 1D**). Thus, culture in F12 increases the susceptibility to several ferroptosis-inducing agents.

### 3.2 Culture in F12 lowers the threshold for iron dependent lipid peroxidation

Since glutathione is required for reduction of lipid hydroperoxides to lipid alcohols by GPX4, depletion of glutathione may sensitize cells to lipid peroxidation [9, 29]. Indeed, BSO treatment (100µM, 24 hours) dramatically increased fluorescence emission of the lipid peroxidation sensor BODIPY-C11 in F12-cultured cells, but only marginally in cells that were cultured in RPMI (**Fig. 2A, B**). There was no difference in general ROS levels between BSO-treated F12 and RPMI-cultured cells, as indicated by the CM-H2DCFDA sensor (**Fig. S2**). Moreover, F12-cultured cells were rescued from lethal BSO concentrations by the lipid peroxide scavengers liproxstatin-1, ferrostatin, or α-tocopherol (**Fig. 2C**). Since lipid peroxidation is catalyzed by ferrous iron in Fenton-like reactions [30], F12-cultured cells were BSO-treated in the presence of the iron chelator deferoxamine. Deferoxamine reduced BSO sensitivity of A549 and H838 cells (**Fig. 2D**).

**Fig. 2.**
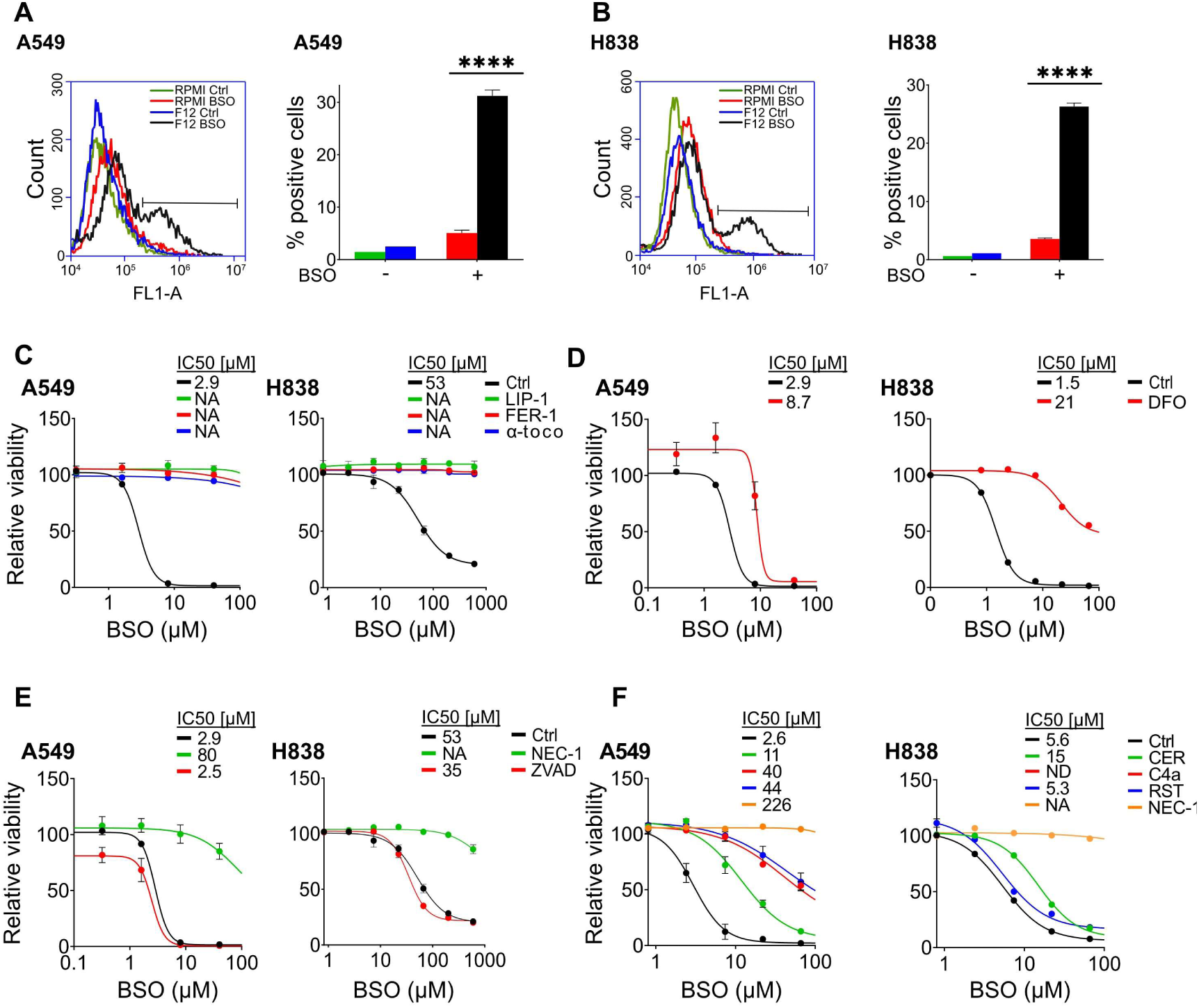
F12 medium sensitizes lung cancer cells to ferroptosis. (**A, B**) FACS count and quantification of BODIPY-C11 fluorescence in A549 (A) or H838 (B) cells that were cultured in RPMI or F12 and treated with 100μM BSO or vehicle for 24 hours. (**C**) Dose response curves for F12-cultured A549 and H838 cells treated with BSO in combination with 5μM liproxstatin-1 (LIP-1), 5μM ferrostatin-1 (FER-1) or 50μM α-tocopherol (α-toco) for 72 hours. The data were normalized against the mean of the untreated samples for each condition. (**D**) Dose response curves for F12-cultured A549 or H838 cells treated with BSO in combination with 5μM deferoxamine (DFO) for A549 cells and 9μM for H838 cells for 72 hours. The data were normalized against the mean of the untreated samples for each condition. (**E**) Dose response curves for F12-cultured A549 or H838 cells treated with BSO in combination with 50μM necrostatin-1 (NEC-1) or 10μM ZVAD-FMK (ZVAD) for 72 hours. The data were normalized against the mean of the untreated samples for each condition. (**F**) Dose response curves for F12-cultured A549 or H838 cells treated with BSO in combination with 4µg/ml certolizumab (CER), 15µM CUCPT4a (C4a), 3μM resatorvid (RST), or 50μM necrostatin-1 (NEC-1) for 72 hours (ND, not done). The data were normalized against the mean of the untreated samples for each condition. n=3 replicates for all datapoints, error bars show SEM. ****P<0.0001

Iron-dependent lipid peroxidation is indicative of ferroptosis but also causes other forms of regulated cell death. Cell death mechanisms are distinguished by their use of distinct effector proteins: caspases in apoptosis and pyroptosis, receptor-interacting serine/threonine protein kinase 1 (RIPK1) in necroptosis, and none of these in ferroptosis [31]. BSO-treated A549 or H838 cells that were cultured in F12 medium were rescued in the presence of the RIPK1 inhibitor necrostatin-1 but not in the presence of the pan-caspase inhibitor Z-VAD-FMK (**Fig. 2E**), suggesting that necroptosis might play a role.

RIPK1 operates downstream of tumor necrosis factor receptor-1 (TNFR1) or toll-like receptor (TLR) 3 or 4 in necroptosis [31]. In line with the necrostatin-1 results, BSO-treated A549 cells were rescued in the presence of the TNFR1 inhibitor certolizumab, the TLR3 inhibitor CU-CPT 4a, or the TLR4 inhibitor resatorvid (**Fig. 2F**). BSO-treated H838 cells were similarly rescued in the presence of certolizumab (**Fig. 2F**), supporting the involvement of necroptosis. The cells were not rescued by resatorvid, and CU-CPT 4a was toxic and therefore not tested, indicating some variation between the cell lines. Overall, the results suggest that F12-culture lowers the threshold for necroptosis in response to BSO-induced iron dependent lipid peroxidation.

### 3.3 The sensitizing effect of F12 is caused by lower amino acid content

To identify which nutritional factors increase sensitivity to BSO in F12-cultured cells, we compared the composition of F12 and RPMI medium. F12 contains less amino acids compared to RPMI, but a wider variety of other components (**Table 1**). To test whether BSO sensitivity is governed by the amount of amino acids, we supplemented F12 medium with 5 times the amount of essential and non-essential amino acids (F12 EXAA in **Table 1**) and established BSO dose response curves. Addition of extra amino acids rescued A549 cells from BSO (**Fig. 3A**).

**Fig. 3.**
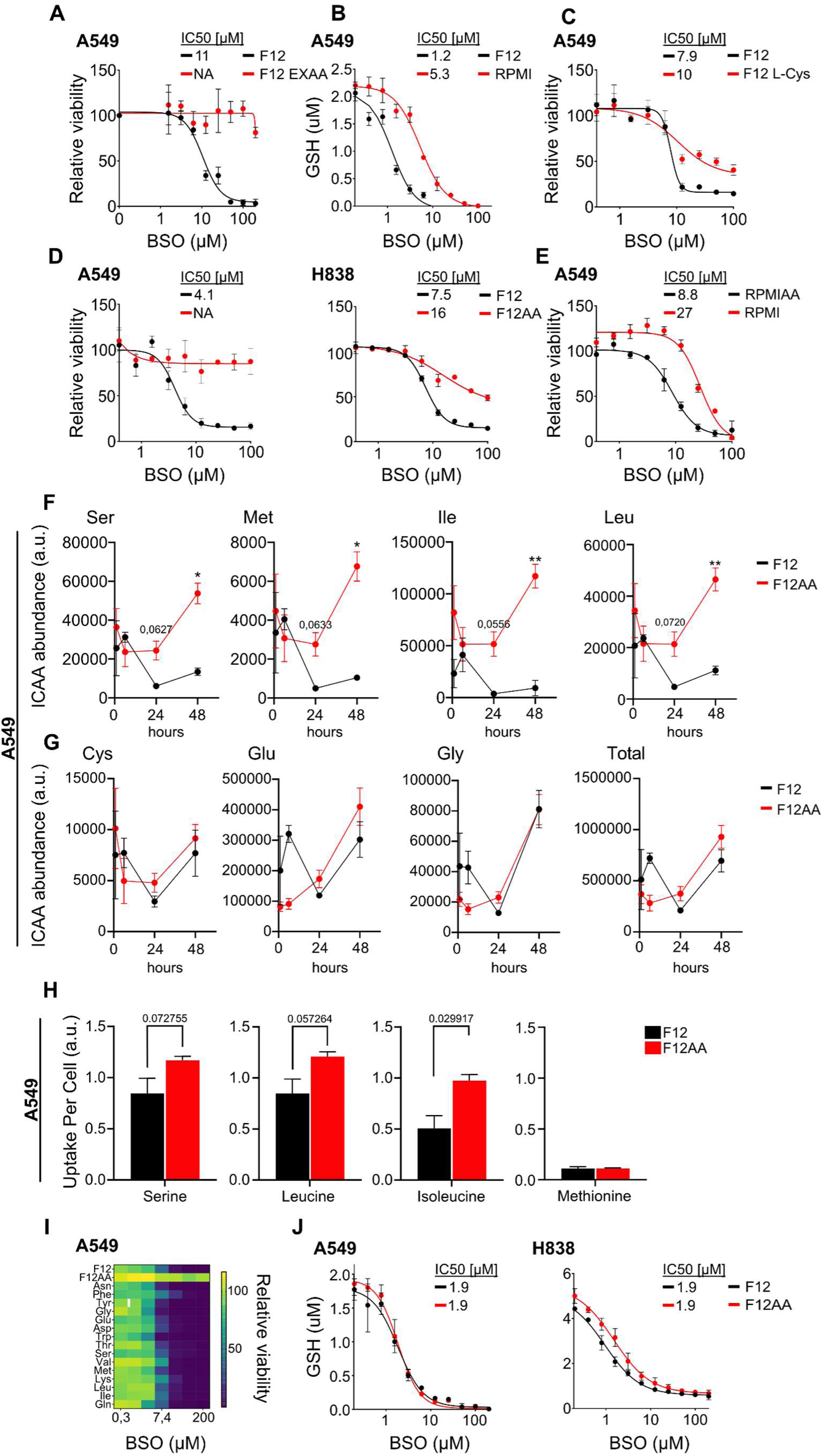
The sensitizing effect of F12 is caused by lower amino acid content. (**A**) BSO dose response curves for A549 cells cultured in F12 or F12 supplemented with extra amino acids (F12EXAA, see Table 1 for media composition) for 72 hours. The data were normalized against the mean of the untreated samples for each condition. (**B**) Concentrations of reduced glutathione in A549 cells cultured in RPMI or F12 and treated with the indicated concentrations of BSO for 24 hours. (**C**) BSO dose response curves for A549 cells cultured in F12 or F12 supplemented with 65mg/L cystine (F12 L-Cys) for 72 hours. The data were normalized against the mean of the untreated samples for each condition. (**D**) BSO dose response curves for A549 and H838 cells cultured in F12 or F12AA for 72 hours. The data were normalized against the mean of the untreated samples for each condition. **(E)** BSO dose response curves for A549 cells cultured in RPMI or RPMIAA, the latter with amino acid concentrations matching those of F12 (see Table 1), for 72 hours. The data were normalized against the mean of the untreated samples for each condition. (**F, G**) GC/MS data for intracellular levels of serine, methionine, isoleucine, and leucine (F) or cysteine, glutamate, and glycine (G) in A549 cells at 1, 6, 24, and 48 hours after switching from RPMI medium to F12 or F12AA medium. **(H)** GC/MS data showing uptake of serine, leucine, isoleucine, and methionine in A549 cells that were cultured in F12 or F12AA medium for 48 hours. **(I)** Heatmap showing BSO dose responses of A549 cells cultured in F12 supplemented with the indicated amino acids at final concentrations matching the ones in RPMI (see Table 1 for media composition). **(J)** Concentrations of reduced glutathione in A549 and H838 cells cultured in F12 or F12AA and treated with the indicated concentrations of BSO for 24 hours. n=3 replicates for all datapoints, error bars show SEM. **P<0.01, *P<0.05

The mass concentration of cystine (a dimer of oxidized cysteine) in RPMI exceeds the mass concentration of cysteine in F12. Since cystine is converted to cysteine in the cytoplasm and availability of cysteine limits glutathione production [24, 32], we expected glutathione concentration and BSO tolerance to be higher in RPMI compared to F12-cultured cells. In agreement, concentrations of glutathione were higher in RPMI-cultured cells after titration with BSO (**Fig. 3B**), and addition of cystine increased the BSO tolerance of F12-cultured cells (**Fig. 3C**).

To test whether addition of amino acids other than cystine alters BSO sensitivity, we used a custom-made F12 medium (F12AA) with amino acid concentrations matching the ones in RPMI, with two modifications: We replaced cystine with cysteine at the same concentration as in F12 to minimize the impact on glutathione production. We also maintained the concentration of glutamine as in F12 since glutamine can affect ferroptosis susceptibility [33, 34]. The formulations of the media used are shown in **Table 1**. The higher concentration of amino acids in F12AA conferred marked resistance to BSO in A549 cells, compared to F12-cultured cells (**Fig. 3D**). Similar results were obtained for H838 cells (**Fig. 3D**). Conversely, A549 cells cultured in a custom-made RPMI medium with amino acid concentrations as in F12 (RPMIAA) were more sensitive to BSO compared to RPMI-cultured cells (**Fig. 3E**).

To assess the media effect on amino acid levels, we determined the amounts of intracellular amino acids in RPMI-cultured A549 cells at different time points after feeding the cells with fresh medium, using gas chromatography-mass spectrometry (GC/MS). With fresh RPMI medium, the concentration of most amino acids increased with a peak at 24 hours before returning toward baseline (**Fig. S3**). In contrast, with F12 medium, amino acid concentrations decreased with a lowest point at 24 hours, raising the possibility that intracellular amino acid depletion contributes to ferroptosis sensitivity (**Fig. S3**). However, levels of intracellular amino acids differed between cells that were given RPMI and F12AA medium that have similar amino acid profiles (**Fig. S3**), showing that media components other than amino acids have a major impact.

To pinpoint effects caused by the amino acid composition of the media, we compared intracellular amino acid concentrations in RPMI-cultured cells receiving fresh F12 or F12AA medium that differ only in amino acid composition. We hypothesized that critical amino acids (1) are depleted over time in F12-cultured cells and (2) are less abundant in F12 compared to F12AA-cultured cells. Four amino acids met these criteria: Levels of serine, methionine, leucine, and isoleucine decreased over time in F12-cultured cells and were lower in F12 compared to F12AA-cultured cells (**Fig. 3F**). The levels of cysteine, glycine and glutamate, which are precursors of glutathione, were not lower in F12 compared to F12AA-cultured cells (**Fig. 3G**), suggesting that the prerequisites for glutathione biosynthesis are not affected. Intracellular levels of all amino acids are shown in (**Fig. S3**).

We then examined amino acid release and uptake by comparing amino acid levels in fresh and conditioned media (**Fig. S4A**). The highest uptake was recorded for serine, leucine, and isoleucine, indicating high consumption of these essential amino acids (**Fig. 3H**). The uptake was lower in F12 compared to F12AA-cultured cells, suggesting that limited availability contributes to the intracellular depletion of these amino acids. There was no difference in the uptake of methionine.

To test whether BSO sensitivity is governed by the media concentration of single amino acids, we supplemented F12 with one amino acid at a time to concentrations matching the ones in RPMI, and established BSO dose response curves for A549 cells. In contrast to extra cystine that increased BSO resistance (**Fig. 3C**), none of the other amino acids including serine, methionine, leucine, and isoleucine substantially affected BSO sensitivity (**Fig. 3I** and **Fig. S4B**). Thus, BSO sensitivity is not governed by the concentration of single amino acids.

Next, we investigated whether the difference in BSO sensitivity between F12 and F12AA cultured cells reflected the availability of glutathione. There was no difference in glutathione concentration between F12- and F12AA-cultured A549 cells, and only a minor difference in H838 cells (**Fig. 3J**). Thus, F12AA-cultured cells remained healthy at BSO concentrations that virtually depleted the cells of glutathione. We conclude that higher concentration of amino acids confers resistance to BSO, not by increasing glutathione availability but by rendering cells independent of glutathione.

The BSO-sensitizing effect of F12 was not observed with all batches of FBS (**Fig. S5A**). With one batch, A549 cells proliferated for several passages in the presence of BSO before they died (**Fig. S5B**). The number of passages that passed before the cells died inversely correlated to the concentration of BSO. This result is reminiscent of a previous study in which BSO induced proteotoxic stress after prolonged treatment [12]. We did not investigate this phenomenon further.

### 3.4 Reduced amino acid levels activates the integrated stress response pathway

Amino acids are used in mRNA translation, as precursors in biosynthesis, as substrates for energy metabolism, and as signaling molecules. To assess whether amino acid signaling plays a role, we investigated the major amino acid sensing pathways: mammalian target of rapamycin complex 1 (mTORC1) – which is particularly sensitive to leucine and isoleucine – and general control nonderepressible-2 (GCN2).

mTORC1 is inactivated at low amino acid concentrations [38]. To assess mTORC1 activity, we investigated phosphorylation of the downstream targets ribosomal protein S6 (S6) and eukaryotic translation initiation factor 4E-binding protein 1 (4E-BP1). There was no decrease in phosphorylation of these proteins in A549 cells that were cultured in F12 versus F12AA or RPMI, in the presence or absence of BSO (**Fig. 4A**). However, total levels of 4E-BP1 were higher in F12 compared to F12AA or RPMI-cultured cells.

**Fig. 4.**
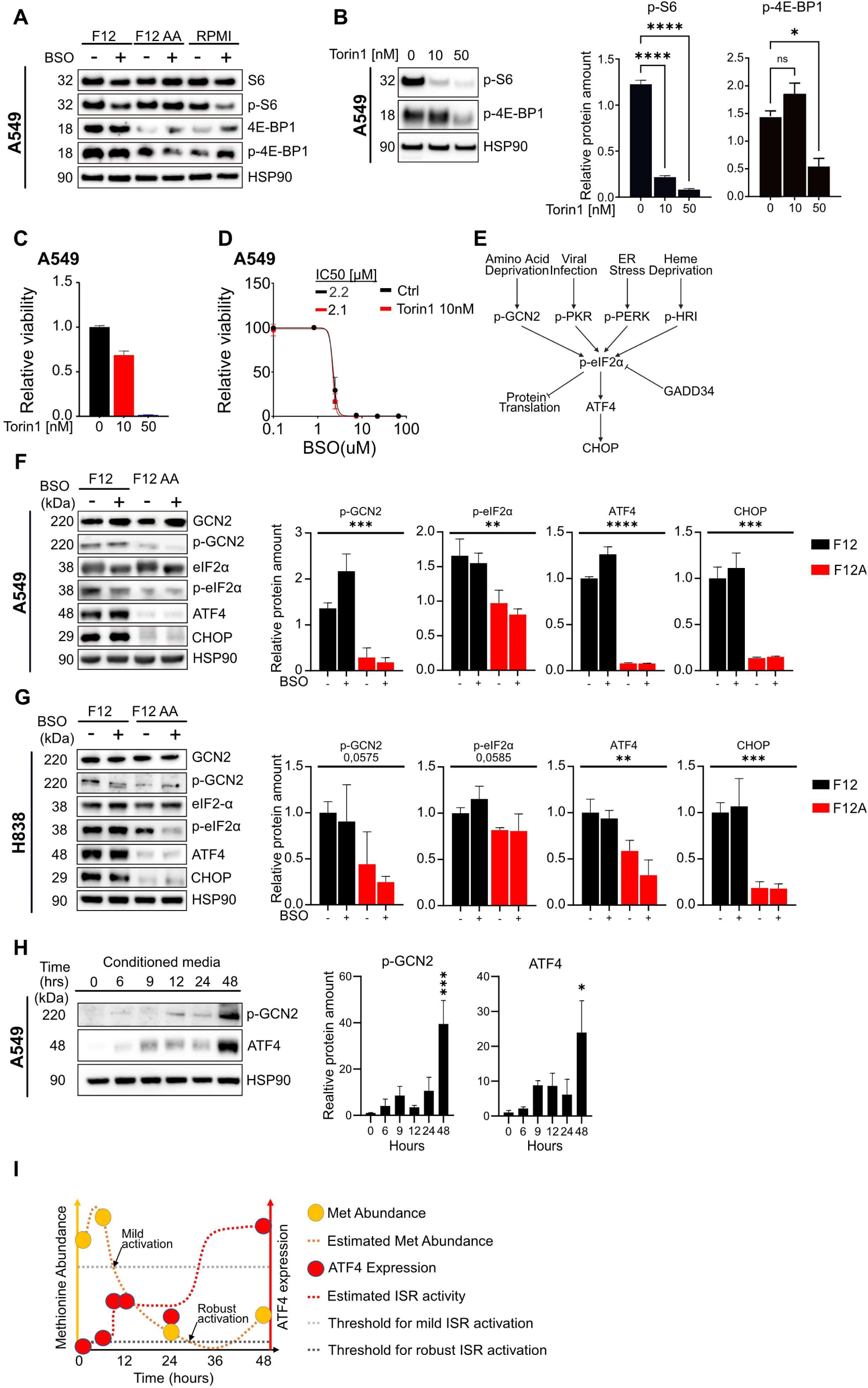
The integrated stress response pathway is activated in F12-cultured cells. (**A**) Western blotting of S6, p-S6, 4E-BP1, p-4E-BP1 in protein extracts of A549 cells cultured in F12, F12AA, or RPMI and treated with 100μM BSO for 24 hours. HSP90 was used as loading control. **(B)** Western blotting and quantification of p-S6 and p-4E-BP1 in protein extracts of A549 cells cultured in F12 and treated with 10 or 50nM torin1 for 24 hours. HSP90 was used as loading control. (**C**) Viability (luminescence) of A549 cells treated with 10 or 50nM torin1 for 24 hours. (**D**) BSO dose response curves for A549 cells cultured in F12 and treated with 10nM torin1 for 72 hours. The data were normalized against the mean of the untreated samples for each condition. **(E)** Schematic of the ISR pathway. (**F, G**) Western blotting and quantification of GCN2, p-GCN2, eIF2α, p-eIF2α, ATF4, and CHOP in protein extracts of A549 (F) or H838 (G) cells cultured in F12 or F12AA and treated with 100μM BSO for 24 hours. HSP90 was used as loading control. **(H)** Western blotting and quantification of p-GCN2 and ATF4 in protein extracts of A549 cells at 0, 6, 9, 12, 24, and 48 hours after switching from F12AA medium to a pre-conditioned F12 medium. (**I**) Schematic model showing methionine abundance, estimated methionine abundance, ATF4 expression, and estimated ISR activity, as indicated. Data on methionine abundance were retrieved from Fig. 3F and ATF4 expression from Fig. 4H. Thresholds for mild and robust ISR activation are indicated by arrows. n=3 replicates for all datapoints, error bars show SEM. ****P<0.0001, ***P<0.001, **P<0.01, *P<0.05

To fully exclude the role of mTORC1, we treated A549 cells with increasing concentrations of the mTOR inhibitor torin1 and determined levels of phosphorylated S6 and 4E-BP1 (**Fig 4B**). Because torin1 decreased cell viability on its own (**Fig. 4C**), we could not use high concentrations of torin1 in combination with BSO. Nevertheless, at concentrations that clearly abolished phosphorylation of S6, torin1 had no impact on the IC50 value of BSO (**Fig 4D**). Thus, inactivation of mTORC1 does not play a role in this scenario.

The GCN2 kinase is activated by auto-phosphorylation in the presence of uncharged tRNA molecules that accumulate when amino acids become sparse [35, 36]. This leads to phosphorylation of eukaryotic translation initiation factor 2α (eIF2α) and activation of the integrated stress response (ISR) pathway (**Fig. 4E**). Phosphorylated eIF2α inhibits CAP dependent mRNA translation while stimulating CAP-independent translation of selected proteins including activating transcription factor 4 (ATF4) and C/EBP homologous protein (CHOP, DDIT3), thus orchestrating the integrated stress response [36, 37].

Phosphorylation of GCN2 and eIF2α was increased in F12 compared to F12AA-cultured A549 and H838 cells, indicating activation of the ISR pathway (**Fig. 4F, G**). In agreement, the amount of ATF4 and CHOP was dramatically increased in F12 compared to F12AA-cultured cells (**Fig. 4F, G**). The expression of these proteins was not affected by BSO treatment, showing that ISR-activation is mediated by low amino acid levels and not by oxidative stress. Activation of the ISR pathway can explain why 4E-BP1, which is an established target of ATF4 [38], showed increased expression in F12 compared to F12AA or RPMI-cultured cells.

eIF2α is activated by additional kinases: RNA-like ER kinase (PERK) in response to endoplasmic reticulum stress, protein kinase R (PKR) in response to virus infection, and eIF2α kinase heme-regulated inhibitor (HRI) in response to heme deprivation (**Fig. 4E**) [36]. PERK and PKR were phosphorylated in A549 and H838 cells and may thus contribute to ISR activity (**Fig. S6A, B**). However, there was no difference in phosphorylation of the proteins between cells cultured in F12 and F12AA, suggesting that they do not trigger activation of the pathway in F12-cultured cells. HRI was not investigated since it’s mainly expressed in erythroid cells [39].

To determine how long it takes to activate the ISR pathway after reducing the amino acid concentration in the medium, we measured ATF4 expression in A549 cells at different time points after switching from F12AA to F12 medium. ATF4 was not upregulated until 48 hours after the switch (**Fig. S6C**) suggesting that amino acid consumption by the cells over time is necessary to reach the threshold concentration in the medium that activates the pathway, or that activation is delayed by cell-intrinsic factors. To distinguish between these possibilities, we repeated the experiment using F12 medium that had been preconditioned with A549 cells for 48 hours and included phospho-GCN2 in the analysis. Induction of GCN2 phosphorylation and ATF4 expression was evident at 9 h but not fully developed until 48 h after the media change (**Fig. 4H**). Thus, activation of the pathway is delayed by cell-intrinsic factors, possibly depletion of intracellular amino acids. Based on our data on methionine abundance (**Fig. 3F**) and ATF4 expression (**Fig. 4H**), we propose a parsimonious scenario for GCN2 activation of the ISR-pathway (**Fig. 4I**): Intracellular concentrations of critical amino acids decline after 6 hours to reach a lowest point between 24 and 48 hours after media exchange. The ISR pathway is mildly activated between 6-9 hours and robustly activated after 24 hours leading to metabolic adaptations that partly restore amino acid concentrations at 48 hours after media exchange.

In a previous publication, the RIPK1 inhibitor necrostatin-1 abolished induction of ATF4-expression in response to cystine starvation in triple-negative breast cancer cells [40], raising the possibility that necrostatin-1 rescued BSO-treated F12-cultured cells (see **Fig. 2E**) by blocking ATF4-expression, and not by inhibiting necroptosis. To exclude this possibility, F12-cultured A549 cells were treated for 24 hours with necrostatin-1. The treatment had no effect on ATF4-expression in F12-cultured cells (**Fig. S6D**). Similar results were obtained with inhibitors of TNFR1, TLR3, or TLR4 that operate upstream of RIPK1 in necroptosis (**Fig. S6D**).

### 3.5 Increased autophagy in F12-cultured cells does not influence BSO sensitivity

Activation of the ISR pathway increases autophagy by which cells degrade their own organelles and macromolecules to recycle metabolites [41]. Since autophagy and ferroptosis are causally interconnected [42], autophagy may contribute to greater BSO sensitivity in F12-cultured cells. We used the lipidated isoform of microtubule-associated protein 1A/1B-light chain 3B (LC3B-II) to monitor the amount of autophagosomes, and the lysosome inhibitor chloroquine to estimate autophagic flux. Following addition of chloroquine, levels of LC3B-II increased at a linear rate during 4 hours, after which the slope levelled off (**Fig. 5A, B**). During the linear phase, the accumulation of LC3B-II was faster in F12 (0.18 units/hr [0.14-0.22]) compared to F12AA-cultured cells (0.12 units/hr [0.09-0.14]), indicating a small but significant increase in autophagic flux.

**Fig. 5.**
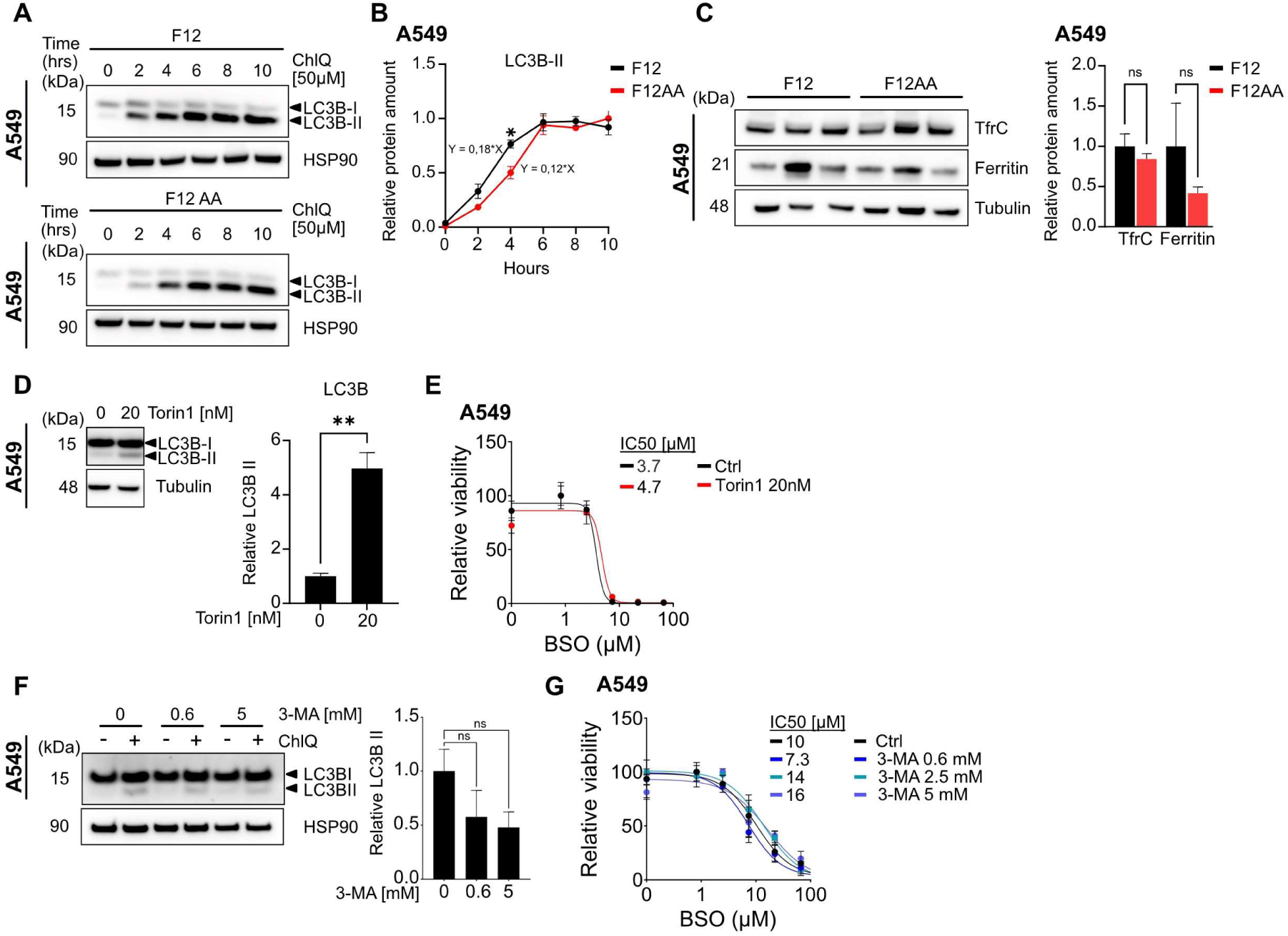
Increased autophagy in F12-cultured cells does not influence BSO sensitivity. (**A, B**) Western blotting (A) and quantification (B) of LC3B-I and II expression in protein extracts of A549 cells cultured in F12 or F12AA and treated with 50μM chloroquine (ChlQ) for the indicated time periods. HSP90 was used as loading control. **(C)** Western blotting and quantification of TFRC and Ferritin (heavy chain) in protein extracts of A549 cells cultured in F12 or F12AA. Tubulin was used as loading control **(D)** Western blotting and quantification of LC3B-II in protein extracts of A549 cells cultured in F12 and treated with 20nM Torin1 for 24 hours. Tubulin was used as loading control. **(E)** Dose response curves for F12-cultured A549 cells treated with BSO in combination with 20nM Torin1 for 72 hours. The data were normalized against the mean of the untreated samples for each condition. (**F**) Western blotting and quantification of LC3B-II expression in protein extracts of A549 cells cultured in F12 and treated with 0.6, 2.5, or 5mM 3-MA in the presence of 50μM chloroquine (ChlQ) for 1 hour. HSP90 was used as loading control. **(G)** Dose response curves for F12-cultured A549 cells treated with BSO in combination with 0.6, 2.5, or 5mM 3-MA for 72 hours. The data were normalized against the mean of the untreated samples for each condition. n=3 replicates for all datapoints, error bars show SEM. **P<0.01, *P<0.05

Ferritinophagy, by which cells recycle ferritin, is particularly relevant to ferroptosis because it increases the labile iron pool [43, 44]. Although ferritin levels varied between samples, there was no evidence of decreased ferritin levels in F12 compared to F12AA-cultured cells (**Fig. 5C**), arguing against increased ferritiophagy as a contributing factor. Moreover, there was no difference in transferrin receptor-1 (TFRC) expression between F12 and F12AA-cultured cells (**Fig. 5C)**, excluding another possible cause of altered ferroptosis susceptibility.

To assess the significance of increased autophagic flux, we treated F12-cultured A549 cells with BSO in the presence of chloroquine or bafilomycin, another lysosome inhibitor. Both inhibitors reduced BSO sensitivity suggesting that BSO-lethality may be autophagy-dependent (**Fig. S7A**). However, prolonged chloroquine treatment (24 h) shut down expression of ATF4 in F12-cultured cells making the data difficult to interpret (**Fig. S7B**). To clarify the importance of autophagy, we used the mTOR inhibitor torin1 to stimulate macroautophagy. Concentrations of torin1 that increased the amount of autophagosomes, as shown by augmented LC3B-II expression (**Fig. 5D**), had no effect on BSO sensitivity (**Fig. 5E**). Moreover, the macroautophagy inhibitor 3-methyladenine (3MA), which reduced accumulation of LC3B-II in the presence of chloroquine, did not alter the BSO IC50 values (**Fig. 5F, G**). Overall, we found no compelling evidence that autophagy affects BSO sensitivity in F12-cultured cells. The torin1 data support our previous assessment that mTORC1 is not involved (see **Fig. 4B-D**).

### 3.6 Activation of the ISR pathway sensitizes lung cancer cells to BSO

The ISR is mainly an adaptive response that decreases cell stress, but prolonged or excessive activation of the pathway can induce cell death [36]. To investigate whether ISR pathway activation attenuates or induces cell death in the presence of BSO, we inactivated GCN2 in F12-cultured A549 or H838 cells using three independent siRNAs (**Fig. 6A, B**). Knockdown of GCN2 reduced levels of phosphorylated eIF2α, ATF4, and CHOP, compared to control (**Fig. 6B** and **Fig. S8A**), showing that GCN2 is required for ISR pathway activation in F12-cultured cells. Two siRNA markedly reduced cell death in the presence of BSO suggesting that ISR pathway activation is required for BSO-lethality (**Fig. 6C**). One of the siRNAs did not affect BSO sensitivity, possibly reflecting off-target toxicity.

**Fig. 6.**
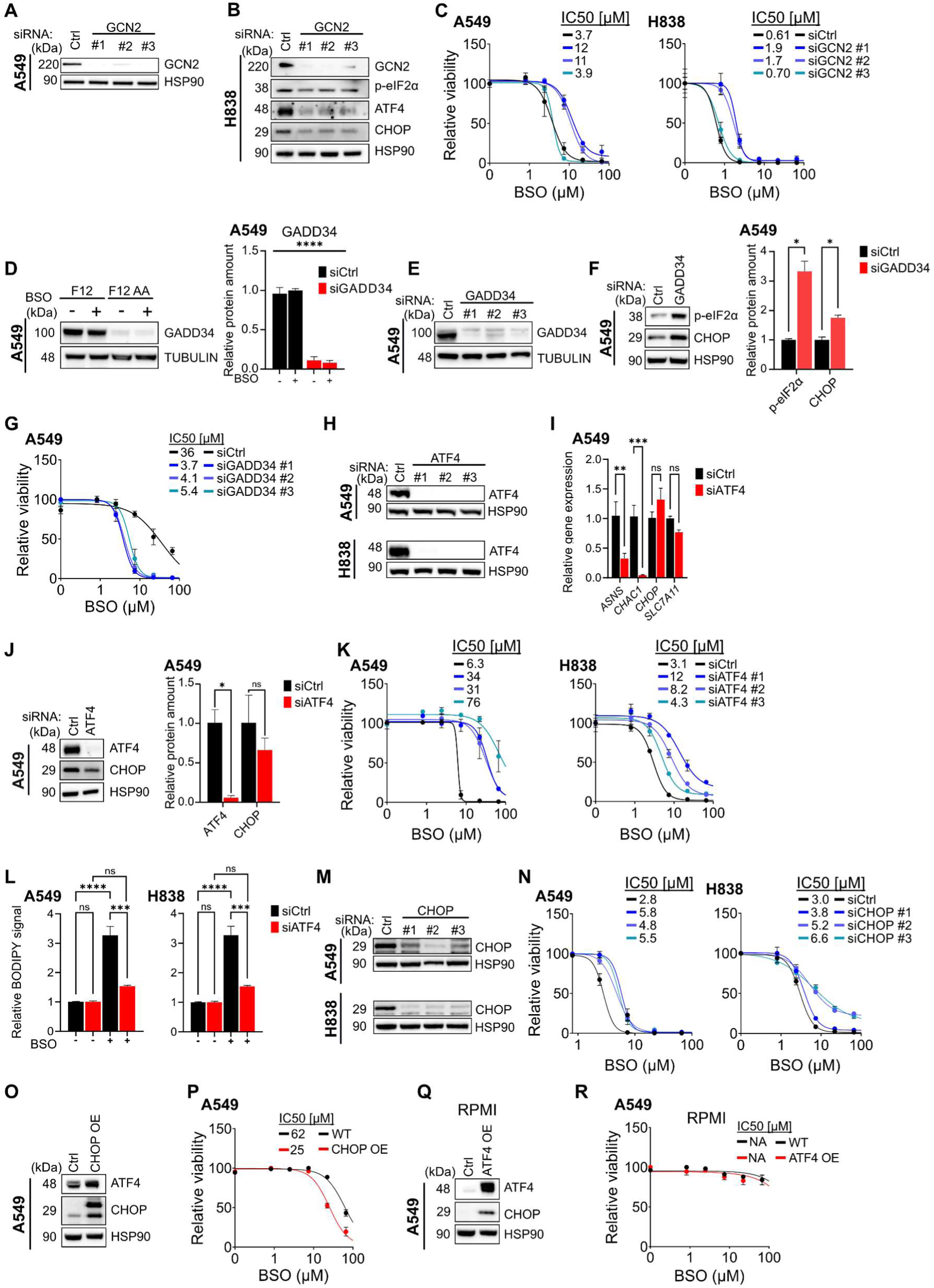
Activation of the ISR pathway sensitizes lung cancer cells to BSO. (**A, B**) Western blotting of GCN2 in protein extracts of A549 cells (A) and GCN2, p-eIF2α, ATF4 and CHOP in protein extracts of H838 cells (B) that were transfected with Ctrl siRNA or siRNA targeting GCN2 mRNA. HSP90 was used as loading control. (**C**) BSO dose response curves for A549 (left) and H838 (right) cells cultured in F12 for 72 hours and transfected with Ctrl siRNA or siRNA targeting GCN2 mRNA. The data were normalized against the mean of the untreated samples for each siRNA. (**D**) Western blotting and quantification of GADD34 in protein extracts of A549 cells that were cultured in F12 or F12AA medium. Tubulin was used as loading control. (**E**) Western blotting of GADD34 in protein extracts of A549 cells that were transfected with Ctrl siRNA or siRNA targeting GADD34 mRNA. Tubulin was used as loading control. (**F**) Western blotting and quantification of p-eIF2α and CHOP in protein extracts of A549 cells that were transfected with Ctrl siRNA or siRNA targeting GADD34 mRNA. HSP90 was used as loading control. (**G**) BSO dose response curves for A549 cells cultured in F12 for 72 hours and transfected with Ctrl siRNA or siRNA targeting GADD34 mRNA. The data were normalized against the mean of the untreated samples for each condition. (**H**) Western blotting of ATF4 in protein extracts of A549 (top) or H838 (bottom) cells that were transfected with Ctrl siRNA or siRNA targeting ATF4 mRNA. HSP90 was used as loading control. (**I**) mRNA expression of ASNS, CHAC1, CHOP, and SLC7A11 in A549 cells transfected with siRNA targeting ATF4. GAPDH was used as a reference gene for normalization. (**J**) Western blotting and quantification of ATF4 and CHOP in protein extracts of A549 cells that were transfected with Ctrl siRNA or siRNA targeting ATF4 mRNA. HSP90 was used as loading control. (**K**) BSO dose response curves for A549 (left) and H838 (right) cells cultured in F12 for 72 hours and transfected with Ctrl siRNA or siRNA targeting ATF4 mRNA. The data were normalized against the mean of the untreated samples for each siRNA. (**L**) FACS quantification of BODIPY-C11 fluorescence in A549 (left) or H838 (right) cells that were cultured in F12 and treated with 100μM BSO or vehicle for 24 hours and transfected with Ctrl siRNA or siRNA targeting ATF4 mRNA. (**M**) Western blotting of CHOP in protein extracts of A549 (top) or H838 (bottom) cells that were transfected with Ctrl siRNA or siRNA targeting CHOP mRNA. HSP90 was used as loading control. ( **N** ) BSO dose response curves for A549 (left) and H838 (right) cells cultured in F12 for 72 hours and transfected with Ctrl siRNA or siRNA targeting CHOP mRNA. The data were normalized against the mean of the untreated samples for each siRNA. (**O**) Western blotting of ATF4 and CHOP in protein extracts of F12-cultured A549 cells that carried a lentivirus overexpressing CHOP cDNA or control. HSP90 was used as loading control. (**P**) BSO dose response curves for A549 cells that carried lentivirus overexpressing CHOP cDNA or control and were cultured in F12 for 72 hours. The data were normalized against the mean of the untreated samples for each condition. (**Q**) Western blotting of ATF4 and CHOP in protein extracts of RPMI-cultured A549 cells that carried a lentivirus overexpressing ATF4 cDNA or control. HSP90 was used as loading control. **(R)** BSO dose response curves for A549 cells that carried a lentivirus overexpressing ATF4 cDNA or control and were cultured in RPMI for 72 hours. The data were normalized against the mean of the untreated samples for each condition. n=3 replicates for all datapoints, error bars show SEM. ****P<0.0001, ***P<0.001, **P<0.01, *P<0.05

To further assess the impact of ISR-pathway activation, we investigated DNA damage-inducible protein 34 (GADD34, PPP1R15A), a negative feedback regulator of the ISR pathway that dephosphorylates p-eIF2α (see **Fig. 4E**) [45]. Expression of GADD34 was markedly higher in F12 compared to F12-cultured cells (**Fig. 6D** and **Fig. S8B**) and knockdown of GADD34 in F12-cultured cells increased phosphorylation of eIF2α, expression of the downstream effector CHOP, and sensitivity towards BSO (**Fig. 6E-G** and **Fig. S8C**), in line with the GCN2 data.

ATF4 orchestrates a transcriptional response downstream of the ISR pathway. To assess the role of ATF4 driven transcription, we knocked down ATF4 (**Fig. 6**H) and investigated mRNA expression of known ATF4 targets [46–49]. Knockdown of ATF4 reduced expression of ASNS and CHAC1, but had no effect on CHOP or SLC7A11 (**Fig. 6I**), suggesting that ATF4 regulates a subset of known targets in F12-cultured cells. Lack of CHOP regulation was confirmed by western blotting (**Fig. 6J**). Knockdown of ATF4 had two effects on cell viability, one in the absence of BSO and one in the presence. In the absence of BSO, knockdown of ATF4 reduced the number of cells, compared to controls, suggesting that ATF4 adapts cells to the low amino acid content in F12 medium (**Fig. S8D**). At the same time, ATF4-deficient cells were markedly resistant to BSO (**Fig. 6K**) showing that a transcriptional response downstream of ATF4 is required for BSO-toxicity. In accord, knockdown of ATF4 prevented lipid peroxidation in the presence of BSO (**Fig. 6L**).

By degrading glutathione, CHAC1 (glutathione-specific γ-glutamylcyclotransferase 1) increases ferroptosis susceptibility in other models [40, 49–51], suggesting that ATF4 driven CHAC1 expression might render F12-cultured cells BSO sensitive. However, protein levels of CHAC1 were equal in F12 and F12AA-cultured cells (**Fig. S8E**) and knockdown of CHAC1 mRNA with three independent siRNAs had no impact on BSO sensitivity in F12-cultured cells (**Fig. S8F, G**). Thus, CHAC1 is not playing an important role in this model. In agreement, glutathione concentrations were equal in F12 compared to F12AA-cultured cells (**Fig. 3J**).

The transcription factor CHOP can induce cell death in response to excessive activation of the ISR pathway [40]. In the absence of BSO, transfection with siRNA against CHOP reduced the number of H838 cells but had no effect on A549 cells, compared to control (**Fig. 6M, Fig. S8H**). In the presence of BSO, inactivation of CHOP conferred partial BSO resistance in both cell lines (**Fig. 6N**). The data suggest that CHOP contributes to BSO sensitivity but not to the same extent as ATF4. However, the inactivation of CHOP was incomplete making the comparison unreliable. To further assess the importance of CHOP, we overexpressed lentiviral CHOP in F12-cultured A549 cells and established BSO dose response curves. CHOP overexpression had a small but significant effect on BSO sensitivity (**Fig. 6O, P**), confirming that CHOP contributes to BSO sensitivity.

To investigate whether expression of ATF4 and CHOP is sufficient to induce BSO sensitivity in RPMI-cultured cells, we infected A549 cells with lentivirus expressing ATF4. The transduced cells overexpressed ATF4 and CHOP (**Fig. 6Q**), consistent with CHOP being an established downstream target of ATF4 in many instances [52]. Overexpression of ATF4 and CHOP had no impact on BSO sensitivity in RPMI-cultured cells (**Fig. 6R**), indicating that amino acid deprivation has additional effects necessary for increasing ferroptosis susceptibility.

We conclude that ATF4 and CHOP orchestrate a transcriptional response downstream of the ISR pathway that help cells to cope with the amino acid sparsity in F12 medium, at the expense of increased susceptibility to lipid peroxidation and ferroptosis-inducing agents such as BSO.

### 3.7 Increased mitochondrial respiration renders F12-cultured cells BSO-sensitive

Cysteine-deprivation-induced ferroptosis requires mitochondrial respiration [53, 54]. To investigate whether mitochondrial function contributes to BSO sensitivity, we measured oxygen consumption rates of A549 and H838 cells grown in F12 or F12AA medium using a Seahorse Extracellular Flux Analyzer. To avoid the possibility that culture media with different pH or buffer capacity would affect the results, we replaced F12 and F12AA medium with assay medium (RPMI) during the analysis. F12-cultured cells showed increased basal and maximal oxygen consumption compared to cells grown in F12AA medium (**Fig. 7A, B**). ATP production was also elevated in F12-cultured cells (**Fig. 7A, B**). Moreover, F12-culture increased the extracellular acidification rate compared to F12AA-culture (**Fig. S9A, B**), suggesting that culture in F12 medium shifted the cells to a more energetic state rather than causing a shift from anaerobic to aerobic respiration. Transfection with siRNA against ATF4 markedly decreased basal and maximal oxygen consumption compared to controls, and reduced ATP-production and extracellular acidification (**Fig. 7C, D** and **Fig. S9C, D**). Thus, increased mitochondrial respiration in F12-cultured cells requires ATF4 expression.

**Fig. 7.**
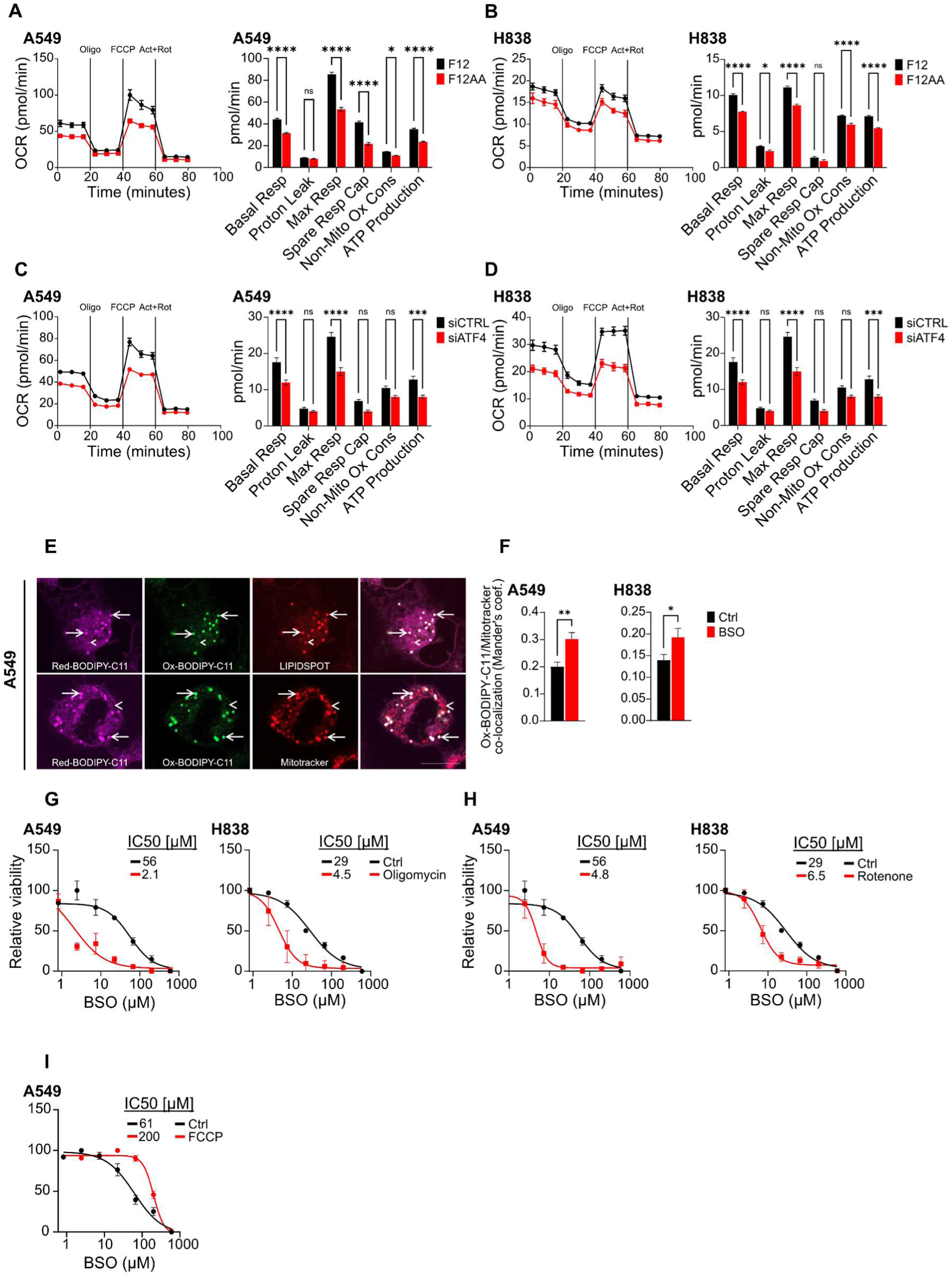
Increased mitochondrial respiration renders F12-cultured cells BSO-sensitive. (**A, B**) Oxygen consumption rate (left) in F12 or F12AA-cultured A549 (A) or H838 (B) cells treated with 0.5μM oligomycin, 1μM FCCP, and 0.5μM rotenone, as indicated. Graphs (right) showing parameter data extracted from the oxygen consumption rate and extracellular acidification rate (n=14 in A549 cells; n=10 in F12 and 4 in F12AA-cultured H838 cells). (**C, D**) Oxygen consumption rate (left) in F12-cultured A549 (C) or H838 (D) cells after transfection with siRNA against ATF4 or control, as above. Graphs (right) showing parameter data, as above (n=15 in A549 cells; n=21 in ATF4 siRNA and 13 in Ctrl siRNA treated H838 cells). (**E**) Confocal microscopy images of F12-cultured A549 cells treated with 100μM BSO for 24 hours showing reduced (pink) and oxidized (green) BODIPY-C11 in combination with lipidspot (red, upper panel) or mitotracker deep red (red, lower panel). Scale bar is 10μm. (**F**) Pixel-wise colocalization (Mander’s coefficient) of oxidized BODIPY-C11 and mitotracker deep red in F12-cultured A549 (left) or H838 (right) cells treated with 100μM BSO for 24 hours or controls (n= 6-9 visual fields in A549 cells and 48 visual fields in H838 cells). (**G-I**) Dose response curves for F12-cultured A549 or H838 cells treated with BSO in combination with 0.5μM oligomycin (G), 0.5μM rotenone (H), or 0.5μM FCCP (I) for 72 hours. The data were normalized against the mean of the untreated samples for each condition. n=3 replicates for all datapoints. Error bars show SEM. ****P<0.0001, ***P<0.001, **P<0.01, *P<0.05

We next examined the spatial relationship between mitochondrial ROS and lipid peroxidation. Using ratiometric confocal microscopy, we found that F12-cultured cells accumulated oxidized BODIPY-C11 within lipid droplets that overlapped with a subset of mitochondria, as evidenced by colocalization with the biomarkers lipidspot and mitotracker deep red, respectively (**Fig. 7E** and **Fig. S9E**). The overlap between oxidized BODIPY-C11 and mitochondria was enhanced by BSO treatment, as shown with pixel-wise colocalization analysis (**Fig. 7F**). Thus, BSO treatment preferentially oxidized BODIPY-C11 in the vicinity of mitochondria. Note that pixel-wise colocalization analysis was performed over entire images and not for regions of interest.

Given that mitochondrial respiration generates superoxide at multiple sites, which is rapidly converted to membrane-permeable hydrogen peroxide that can cause lipid peroxidation [55–57], we next tested whether increasing mitochondrial ROS production could enhance ferroptosis susceptibility. Complex I inhibitors such as rotenone and the ATP synthase inhibitor oligomycin increase superoxide leakage from mitochondria [58–60]. Both drugs markedly increased BSO sensitivity in F12-cultured A549 and H838 cells, indicating that ROS production from mitochondrial respiration is rate-limiting (**Fig. 7G, H**). In contrast, treatment with the uncoupling agent FCCP, which reduces ROS production [61], rescued the viability of BSO-treated A549 cells but had no effect on H838 cells (**Fig. 7I** and **Fig. S9F**). We conclude that culture in F12 medium increases mitochondrial respiration in an ATF4-dependent manner. This increase in respiration promotes ROS production from mitochondria which in tun increases the sensitivity to ferroptosis in F12-cultured cells upon glutathione depletion.

## 4. Discussion

Glutathione is the most abundant intracellular antioxidant, yet glutathione biosynthesis blockade has little impact on the viability of human cultured cancer cells [12–14]. In the present study, we found that levels of extracellular amino acids markedly impact the susceptibility to glutathione depletion in human lung cancer cells. A reduction in the amount of extracellular amino acids activates GCN2 and the ISR pathway, thus sensitizing cancer cells to iron-dependent lipid peroxidation and cell death in the presence of BSO (**Fig. 8**).

**Fig. 8.**
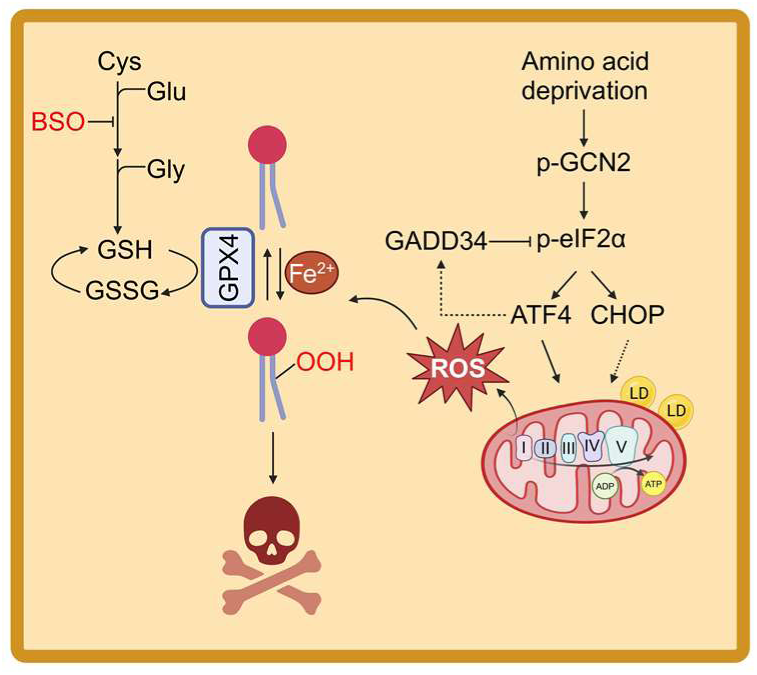
Schematic showing how amino acid restriction sensitizes lung cancer cells to ferroptosis-inducing agents by GCN2-dependent activation of the ISR.

Ferroptosis susceptibility has been previoulsy linked to perturbations limiting glutathione biosynthesis, including cystine or cysteine depletion [62–64], pharmacological blockade of cystine uptake by the system-xc inhibitor erastin [10, 65], or methionine depletion that blocks generation of cysteine through the transsulfuration pathway [19, 66, 67]. The mechanism we describe is uncoupled from cysteine metabolism and glutathione biosynthesis. The concentration of cysteine was equal in F12 and F12AA media, there was no significant difference in intracellular concentrations of cysteine, glycine or glutamate between F12 and F12AA-cultured cells, and there was no difference in glutathione concentration between BSO-treated F12 and F12AA-cultured cells. Instead, ferroptosis susceptibility was linked to glutathione-dependency: At lower concentrations of amino acids, cells become dependent on glutathione to prevent accumulation of lipid peroxides.

Previous publications indicate that activation of the ISR pathway occurs in association with ferroptosis induction by cysteine or cystine depletion, erastin treatment, or genetic inactivation of system-xc, but the significance of this activation has not been thoroughly investigated [40, 65, 68–71]. Our findings shed new light on these observations and on the acknowledged mystery of why depletion of cysteine (or cystine) leads to worse ferroptosis compared to glutathione synthesis blockade alone [12–14, 64, 70–74]: Besides limiting glutathione biosynthesis, cysteine depletion activates the ISR pathway, thus inducing glutathione-dependency.

Activation of the ISR pathway is mediated by four eIF2α kinases; GCN2, PERK, PKR or HRI [36]. Several lines of evidence indicate that GCN2 is the activator in F12-cultured cells: F12-culture induces GCN2 autophosphorylation, knockdown of GCN2 attenuates the expression of ISR markers, and GCN2 knockdown rescues F12-cultured cells in the presence of BSO. PERK and PKR likely feed into the phosphorylation of eIF2α but their activity was not altered by F12-culture. GCN2 is activated by accumulation of uncharged tRNA molecules or by ribosomal stalling or collision [75–77]. We favour a scenario in which GCN2 activation is caused by intracellular depletion of amino acids—specially methionine, serine, leucine, and isoleucine whose levels inversely correlated with GCN2 phosphorylation—and accumulation of the cognate uncharged tRNAs.

The transcription factors ATF4 and CHOP are translated from internal ribosomal entry sequences in response to eIF2α phosphorylation [37, 38]. CHOP, which is a transcriptional target of ATF4 and often positioned downstream of ATF4 in the ISR pathway [52], was regulated independently of ATF4 in F12-cultured cells. Knockdown of ATF4 or CHOP reduced BSO sensitivity, demonstrating that both proteins are required to achieve maximal ferroptosis sensitivity. Since ATF4 and CHOP form heterodimers that co-regulate many genes [78, 79], the proteins may act in concert. However, inactivation of ATF4 had a greater impact on BSO sensitivity than CHOP inactivation, suggesting that ATF4 plays a more important role. Overexpression of ATF4 and CHOP was not sufficient to increase BSO sensitivity in RPMI-cultured cells, demonstrating that F12-culture triggers additional events.

We found that ATF4-expression increases mitochondrial respiration in F12-cultured cells. We propose, based on two lines of evidence, that lipid peroxidation is caused by increased ROS leakage from mitochondria: (1) In the presence of BSO, oxidized BODIPY-C11 accumulated in lipid droplets that colocalize with mitochondria, thus demonstrating spatial correlation between lipid peroxidation and mitochondria. (2) Treatment with rotenone or oligomycin (which increase ROS leakage from the electron transport chain) caused cell death in the presence of BSO, while FCCP treatment (which decreases ROS leakage) made A549-cells less BSO-sensitive. In agreement, ROS generation from mitochondria is an established cause of ferroptosis-susceptibility [11, 53, 80, 81]. ATF4 expression can stimulate or attenuate mitochondrial respiration in different models [82, 83], indicating that ATF4-effects on mitochondrial respiration are controlled by contextual factors that warrant further investigation.

Mitochondria colocalized with lipid droplets containing oxidized lipids in BSO-treated cells. While lipid droplets suppress ferroptosis by sequestering polyunsaturated fatty acids (PUFAs), excessive lipid droplet breakdown can damage cells by releasing oxidized PUFAs [84–87]. Thus, lipophagy mediating lipid droplet breakdown may contribute to ferroptosis susceptibility in F12-cultured cells, a notion supported by the rescue with chloroquine or bafilomycin in BSO-treated cells. However, chloroquine treatment shut down ATF4 expression in F12 cultured cells and ATF4 knockdown rescued the viability of BSO-treated cells, raising the possibility that chloroquine treatment primarily reduces ISR activity. Activation of PERK and the ISR in U2OS cells promotes lipid droplet biogenesis and synthesis of triglycerides containing PUFAs [83]. It is possible that GCN2 activation in F12-cultured cells causes a similar change in lipid metabolism, promoting the formation of lipid droplets prone to oxidation.

BSO is a classic ferroptosis-inducing agent but can also lead to other forms of cell death. Necroptosis is triggered by stimulation of TNFR1 or TLRs and mediated by RIPK1 [31], inhibitors of which rescued BSO-treated cells. Hence, it is possible that necroptosis is part of the killing mechanism. Oxidized phospholipids and their degradation products malondialdehyde or 4-hydroxynonenal serve as damage associated molecular patterns ( DAMPs) for the regulation of innate immunity by pattern recognition receptors such as TLR3 and TLR4 [88]. Moreover, oxidized 1-palmitoyl-2-arachidonoyl-phosphatidylcholine induces lung injury and cytokine production in mice, in a TLR4-dependent manner [89]. Thus, oxidized phospholipids may directly or indirectly stimulate necroptosis-inducing receptors. However, the distinction between ferroptosis and necroptosis is currently vague and should not be overinterpreted [11].

ATF4 can influence ferroptosis susceptibility by preserving cysteine. On one hand, ATF4 stimulates conversion of serine to cysteine through the transsulfuration pathway and cystine import through system-xc. These actions promote glutathione production and protects against ferroptosis [61–67]. On the other hand, ATF4 salvages cysteine from glutathione by upregulating CHAC1, a glutathione-degrading enzyme causing ferroptosis susceptibility [40, 49–51]. Knockdown of ATF4 decreased CHAC1 mRNA, raising the possibility that ATF4-driven CHAC1 expression underpins ferroptosis susceptibility in F12-cultured cells. However, CHAC1 protein levels were equal in F12 and F12AA-cultured A549 cells, CHAC1 siRNAs had no effect on BSO sensitivity, and there was no difference in glutathione concentration between F12 and F12AA-cultured cells. Thus, CHAC1 is not rate limiting for ferroptosis susceptibility in this model. Prolonged methionine deprivation blocks translation of CHAC1 mRNA in human HT-1080 cells and mouse Hepa1–6, MC38, and PanO2 cells, thus uncoupling protein levels from effects of mRNA regulation [51, 66]. It is possible that depletion of methionine in F12-cultured cells has a similar effect.

Nutrient stress is common in tumors due to poor vascularization. Efforts to restore homeostasis by inducing the ISR pathway may collaterally sensitize tumor cells to lipid peroxidation making the cells susceptible to ferroptosis, as shown here. Insufficient amino acid supplies may therefore contribute to the oxidative stress weakness that is evident in many tumors [7, 90–93]. In agreement, serine and glycine restricted diets hampered formation of lymphomas and intestinal tumors in mice, in a ROS-dependent manner [94, 95]. Similarly, cysteine and methionine restricted diets potentiated the effect of ferroptosis-inducing agents on glioma and skin cancer development in mice [51, 66]. Whether a protein restricted diet potentiates the anti-cancer effect of BSO remains to be tested.

Sensitivity to BSO varied between experiments. This could be explained by the fact that Hippo signaling is activated by cell-cell contacts to varying degrees depending on cell density [96], which is a major modifier of BSO-sensitivity in our model. More importantly, the fold change in IC50 values between F12 and F12AA cultured cells was robust as it remained constant across all experiments. Ferroptosis is implicated in vascular and neurodegenerative disorders. Whether amino acid restriction affects ferroptosis susceptibility of non-cancerous cells remains to be tested. Of note, the BSO-sensitizing effect of amino acid restriction was evolutionarily conserved in mouse lung cancer cells (KP cells) and present in SKNBE-2 neuroblastoma cells, indicating that the mechanism might be widespread.

In summary, we show that a mild reduction in levels of extracellular amino acids sensitizes lung cancer cells to ferroptosis-inducing agents by upregulating the ISR pathway. The finding opens new opportunities for dietary amino acid restriction to modify ferroptosis susceptibility in cancer patients.

## Acknowledgements

We acknowledge the Centre for Cellular imaging at the university of Gothenburg and the National Microscopy Infrastructure, NMI (VR-RFI 2019-00217) for providing assistance in microscopy and Malin Lindén for kindly providing the antibody for CHOP. The project was supported by the Swedish Cancer Society (P.L, V.I.S), The Swedish Child Cancer Foundation (P.L.), The Swedish Research Council (P.L., V.I.S, C.W), the Sahlgrenska University Hospital ALF Research (P.L., V.I.S, C.W), and the Sahlgrenska Academy (P.L).

## Notes

### Competing Interest Statement

The authors have declared no competing interest.

