## Supplementary Figures S1-S9 for "Amino acid restriction sensitizes lung cancer cells to ferroptosis via GCN2-dependent activation of the integrated stress response"

### Supplementary Fig. S1

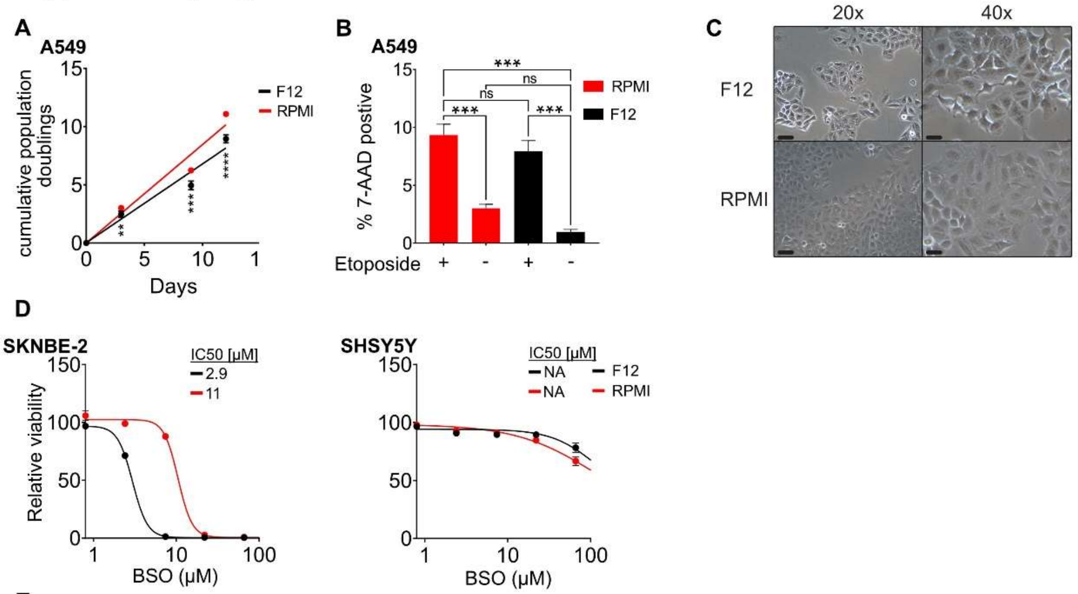


E

| | ★ | ★ | ★ | ★ | | | ★ | ★ |
|---|---|---|---|---|---|---|---|---|
| **Gene** | **A549** | **H838** | **H1299** | **H23** | **H460** | **SHSY5Y** | **SKNBE-2** | **KP** |
| *KRAS* | G12S | - | - | G12C | Q61H | - | - | G12D |
| *ZFHX3* | L1473F | - | F3044V | I1559I | - | - | - | |
| *STK11* | Q37* | T212fs | - | W332* | Q37* | - | - | |
| *TP53* | - | E62* | - | M246I | - | - | C135F | -/- |
| *NRAS* | - | - | Q61K | - | - | - | - | |
| *SMARCA4* | - | - | DEL | - | - | - | - | |
| *PIK3CA* | - | - | - | - | E545K | - | - | |
| *ALK* | - | - | - | - | - | F1174L | - | |
| *NF1* | - | - | - | - | - | - | N664fs | |
| *KEAP1* | G333C | E444 | - | Q193H | D236H | - | - | - |

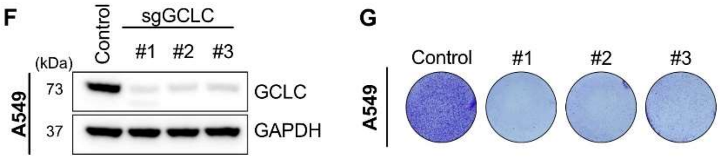


**Fig. S1.** (**A**) Cell growth curves for F12 and RPMI-cultured A549 cells. (**B**) The percentage of 7-AAD-positive A549 cells cultured in F12 and RPMI and treated with etoposide or vehicle for 24 hours. (**C**) Photomicrographs of A549 cells cultured in F12 or RPMI. Scale bars are 60 μm in 20X and 30 μm in 40X. (**D**) BSO dose response curves for SKNBE2 (left) or SHSY5Y (right) neuroblastoma cells cultured in F12 or RPMI for 72 hours. The data were normalized against the mean of the untreated samples for each condition. (**E**) Table showing presence of COSMIC mutations in the investigated cell lines. Stars indicate BSO sensitive cell lines. (**F**) Western blotting of GCLC in protein extracts of A549 cells, 7 days after the cells were infected with lentivirus expressing one of three independent gRNAs targeting coding sequence in GCLC, or non-targeting control gRNA. GAPDH was used as loading control. (**G**) Crystal violet staining of F12-cultured A549 cells, 13 days after the cells were infected with lentivirus expressing gRNAs against GCLC, or non-targeting control gRNA (see methods for details). n=3 replicates for all datapoints, error bars show SEM. ****$P$<0.0001, ***$P$<0.001, **$P$<0.01

### Supplementary Fig. S2

**A**

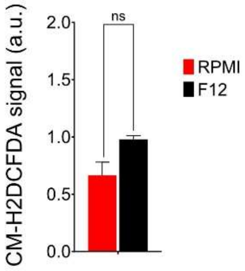


**Fig. S2.** (**A**) CM-H2DCFDA signal in F12 or RPMI-cultured A549 cells treated with 100µM BSO for 24 hours. n=3 replicates, error bars show SEM.

### Supplementary Fig. S3

**A**

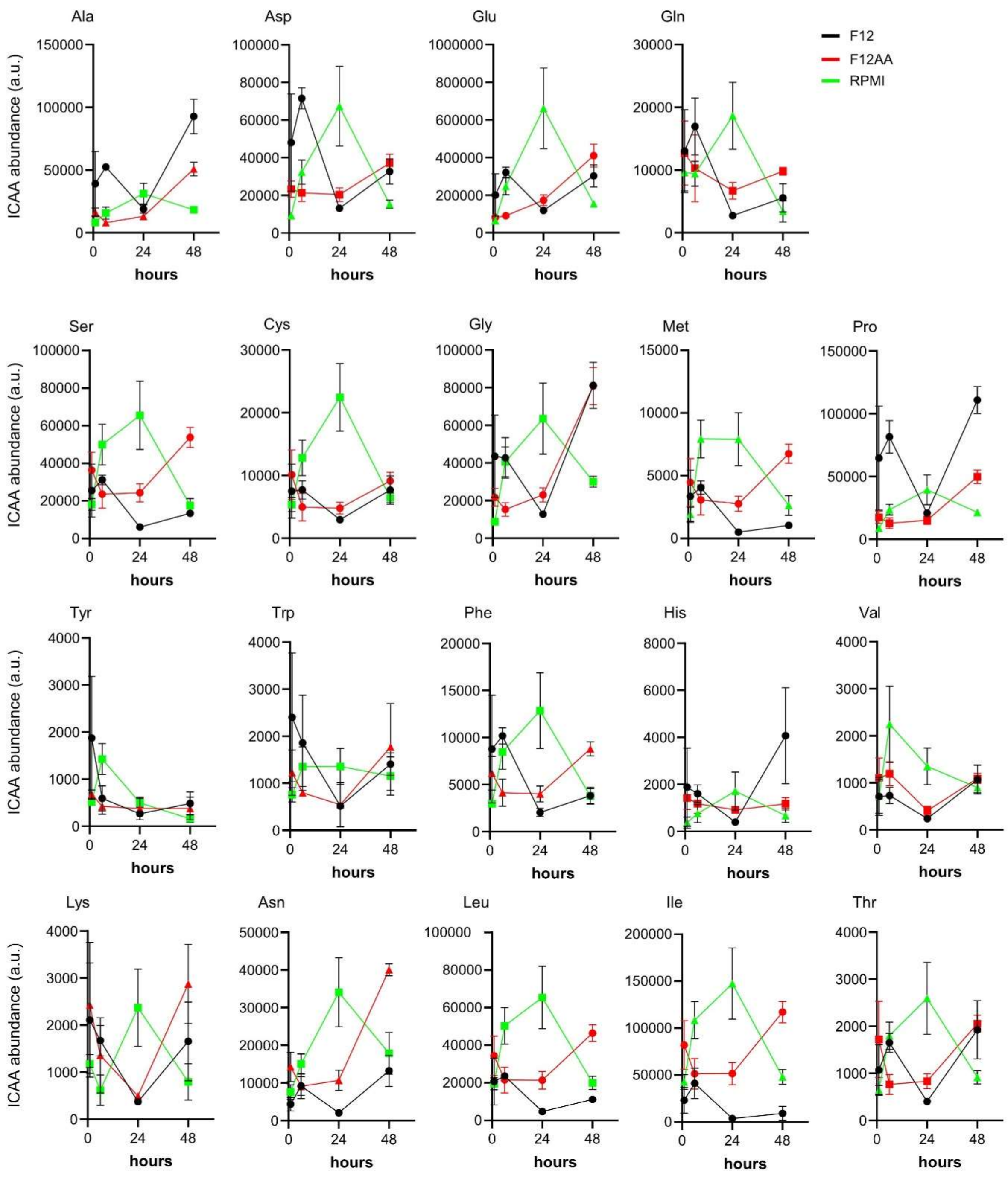


**Fig. S3.** (**A**) GC-MS data for intracellular levels of the indicated amino acid in A549 cells at 1, 6, 24, and 48 hours after switching from RPMI medium to F12, F12AA, or RPMI medium, as indicated. n=3 replicates for all datapoints, error bars show SEM.

### Supplementary Fig. S4

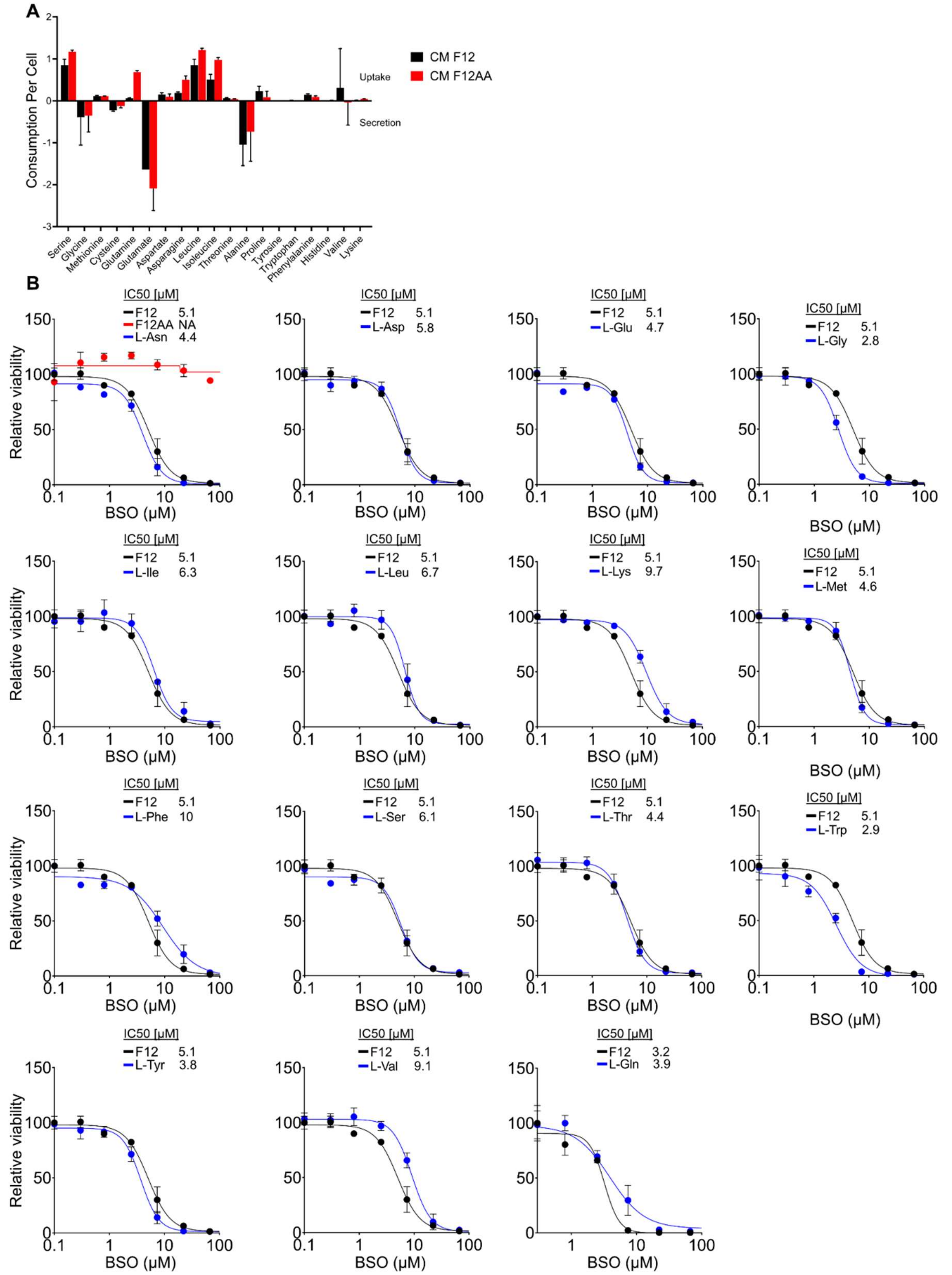


**Fig. S4.** (**A**) GC-MS data showing release (negative values) or uptake (positive values) of the indicated amino acids in A549 cells that were cultured in F12 or F12AA for 48 hours. (**B**) BSO dose response curves for A549 cells cultured in F12 or F12 medium supplemented with single amino acids, as indicated, for 72 hours. The total concentration of the supplemented amino acid was adjusted to match the  concentration in RPMI. The data were normalized against the mean of the untreated samples for each condition. n=3 replicates for all datapoints, error bars show SEM. $**P<0.01$, $*P<0.05$

### Supplementary fig. S5

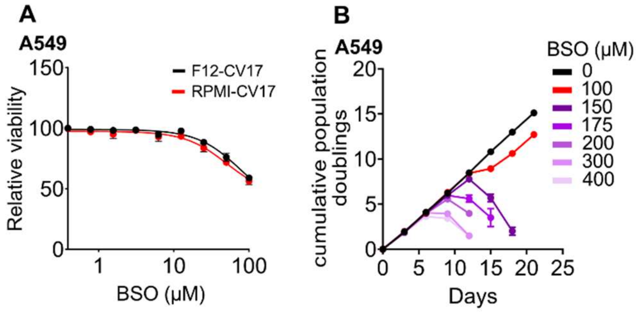


**Fig. S5.** (**A**) BSO dose response curves for A549 cells cultured in F12 or RPMI, supplemented with FBS lot 325079017 (CV17) (Corning), for 72 hours. (**B**) Cell growth curves of A549 cells cultured in F12 supplemented with FBS lot RZJ35921 (HyClone), in the presence of the indicated concentrations of BSO. n=3 replicates for all datapoints, error bars show SEM.

### Supplementary Fig. S6

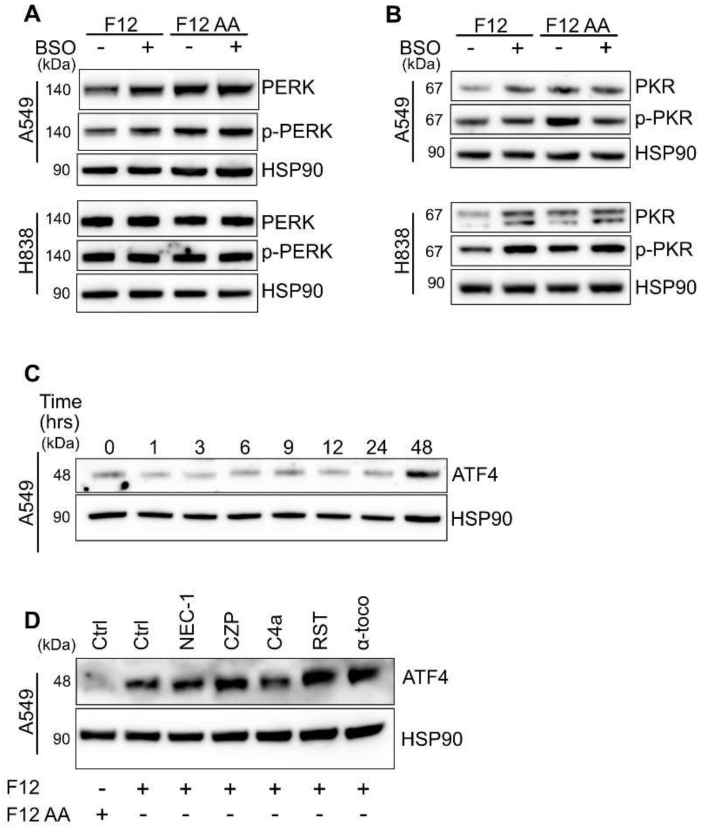


**Fig. S6.** (**A, B**) Western blotting of PERK and p-PERK (A) or PKR and p-PKR (B) in protein extracts of A549 (top) or H838 (bottom) cells cultured in F12 or F12AA, after treatment with 100µM BSO for 24 hours. HSP90 was used as loading control. (**C**) Western blotting of ATF4 in protein extracts of A549 cells at 0, 1, 3, 6, 9, 12, 24, and 48 hours after switching from F12AA medium to F12 medium. (**D**) Western blotting of ATF4 in protein extracts of F12-cultured A549 cells that were treated with necrostatin-1 (NEC-1), certolizumab (CZP), CU-CPT 4a (C4a), resatorvid (RST), or α-tocopherol (α-toco) for 24 hours. HSP90 was used as loading control.

### Supplementary Fig. S7

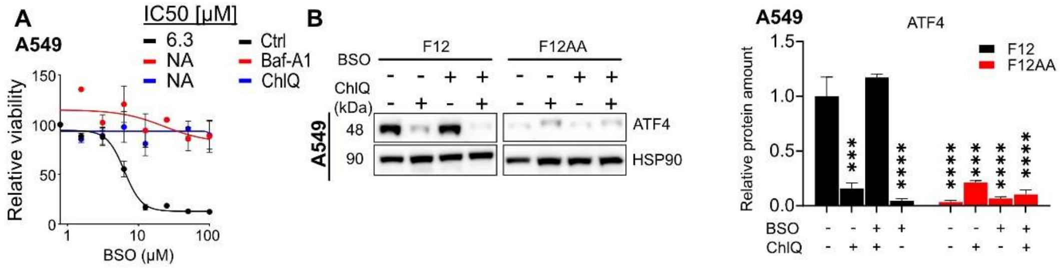


**Fig. S7.** (**A**) Dose response curves for F12-cultured A549 cells treated with BSO in combination with 25µM chloroquine (ChlQ) or 3nM bafilomycin (Baf-A1) for 72 hours. The data were normalized against the mean of the untreated samples for each condition. (**B**) Western blotting and quantification of ATF4 in protein extracts of A549 cells cultured in F12 and treated with 100µM BSO and 50µM ChlQ for 24 hours, as indicated. n=3 replicates for all datapoints, error bars show SEM. ****P<0.0001, ***P<0.001

### Supplementary Fig. S8

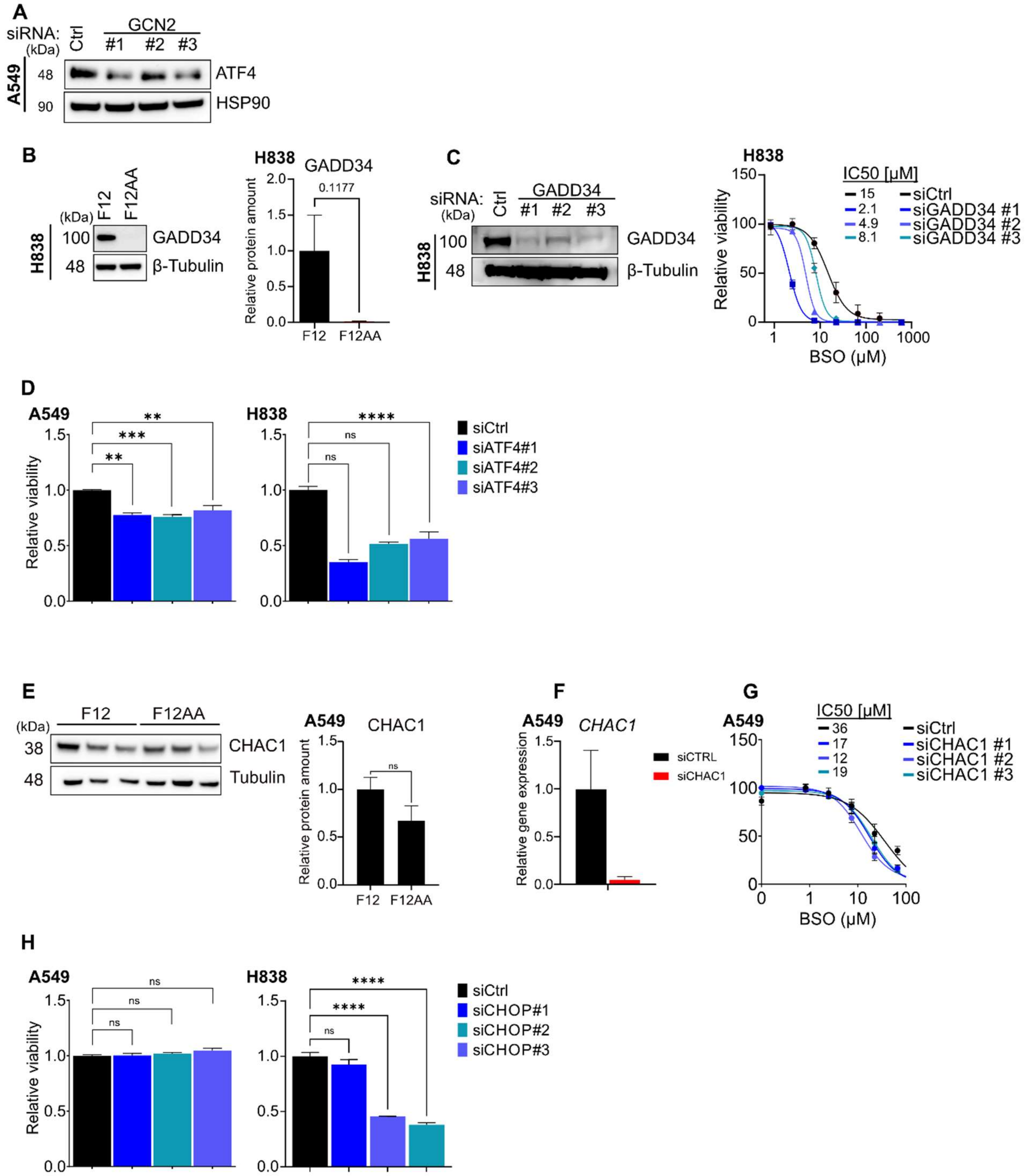


**Fig. S8.** (**A**) Western blotting of ATF4 in protein extracts of F12-cultured A549 cells that were transfected with Ctrl siRNA or siRNA targeting GCN2 mRNA. HSP90 was used as loading control. (**B**) Western blotting and quantification of GADD34 in protein extracts of F12 or F12AA-cultured H838 cells. Tubulin was used as loading control. (**C**) Western blotting (left) of GADD34 in protein extracts of H838 cells that were transfected with Ctrl siRNA or siRNA targeting GADD34 mRNA. Tubulin was used as loading control. BSO dose response curves (right) for H838 cells cultured in F12 for 72 hours and transfected with Ctrl siRNA or siRNA targeting GADD34 mRNA. The data were normalized against the mean of the untreated samples for each siRNA. (**D**) Viability of A549 or H838 cells that were transfected with siRNA against ATF4 and cultured in F12 for 72 hours. Controls were transfected with non-targeting siRNA (Ctrl). (**E**) Western blotting and quantification of CHAC1 in protein extracts of F12 or F12AA-cultured A549 cells. Tubulin was used as loading control. (**F**) mRNA expression of CHAC1 in F12-cultured A549 cells transfected with siRNA targeting CHAC1 mRNA. Controls were transfected with non-targeting siRNA (Ctrl). GAPDH was used as a reference gene for normalization. (**G**) BSO dose response curves for A549 cells cultured in F12 for 72 hours and transfected with siRNA against CHAC1 mRNA. Controls were transfected with non-targeting siRNA (Ctrl). The data were normalized against the mean of the untreated samples for each siRNA. (**H**) Viability of A549 or H838 cells that were transfected with siRNA against CHOP and cultured in F12 for 72 hours. Controls were transfected with non-targeting siRNA (Ctrl). n=3 replicates for all datapoints, error bars show SEM. ****P<0.0001, ***P<0.001, **P<0.01

### Supplementary Fig. S9

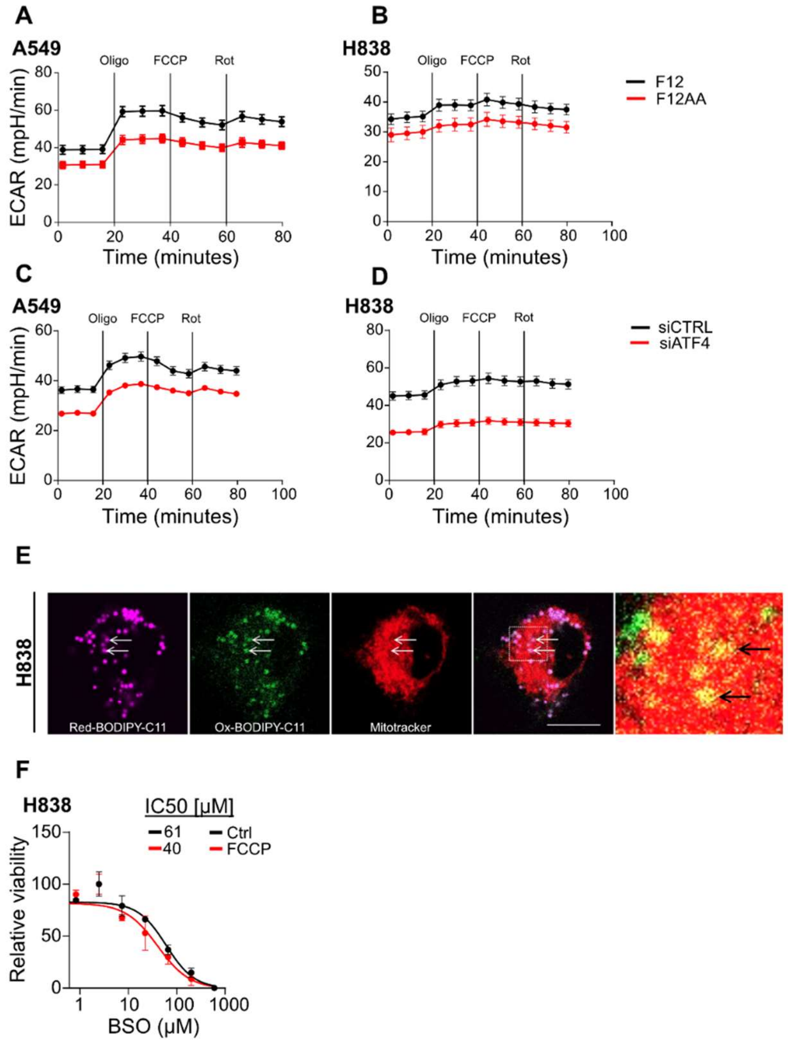


**Fig. S9.** (**A, B**) Extracellular acidification rate (ECAR) in F12 or F12AA-cultured A549 (A) or H838 (B) cells treated with 0.5µM oligomycin, 1µM FCCP, and 0.5µM rotenone, as indicated (n=14 in A549 cells; n=10 in F12 and 4 in F12AA-cultured H838 cells). (**C, D**) ECAR in F12-cultured A549 (C) or H838 (D) cells after transfection with siRNA against ATF4 or control, as above (n=15 in A549 cells; n=21 in ATF4 siRNA and 13 in Ctrl siRNA treated H838 cells). (**E**) Confocal microscopy images of F12-cultured H838 cells treated with 100µM BSO for 24 hours showing reduced (pink) and oxidized (green) BODIPY-C11 in combination with mitotracker deep red (red). Arrows indicate mitochondria-tethered lipid droplets with oxidized BODIPY-C11. The panel at far right shows the dashed area at higher magnification (Scale bar is 10µm). (**F**) Dose response curves for F12-cultured H838 cells treated with BSO in combination with 0.5µM FCCP for 72 hours. The data were normalized against the mean of the untreated samples for each condition (n=3 replicates for all datapoints). Error bars show SEM.
